# coRIC map: a framework for the exploration of multifaceted subcellular RNA-binding proteins

**DOI:** 10.1101/2023.11.06.565240

**Authors:** Yuan Lv, Xiangpeng Guo, Jieyi Hu, Shahzina Kanwal, Jianwen Yuan, Muqddas Tariq, Jingxia Zheng, Mingwei Sun, Yuanyuan Lu, Jingjing Wang, Mengling Jiang, Aiping Wang, Alvaro Castells-Garcia, Xiyuan Zheng, Bin Peng, Dongye Wang, Giacomo Volpe, Liang Wu, Md. Abdul Mazid, Wenjuan Li, Yiwei Lai, Francesca Aguilo, Yu Zhou, Maria Pia Cosma, Xingzhi Xu, Emma Lundberg, Jan Mulder, Andrew P. Hutchins, Patrick H. Maxwell, Luciano Di Croce, Xiaofei Zhang, Miguel A. Esteban

**Author notes:** These authors contributed equally: Yuan Lv, Xiangpeng Guo, Jieyi Hu, Shahzina Kanwal. Corresponding author: Yuan Lv, Xiaofei Zhang, Miguel A. Esteban.

## Abstract

The ability of RNA-binding proteins (RBPs) to form complexes with other biomolecules underpins a broad range of functions and remarkable structural properties. Understanding the precise spatiotemporal dynamics of RBPs and their interacting partners in the steady state and upon perturbation is key to deciphering these aspects. Here, we present the coRIC (<u>co</u>mpartmentalized <u>R</u>NA Interactome <u>C</u>apture) map, an experimental resource and analytical pipeline to study the subcellular dynamics of RBPs through multimodal dataset integration and machine learning. Using this approach, we have generated an atlas comprising 1,768 RBPs distributed in a broad panel of subcellular compartments and delineated their intermolecular and intercompartmental relationships. We have also defined the hierarchy of RBP-containing complexes at multiple scales across the cell, uncovering previously unknown functions of multiple RBPs. Furthermore, we have investigated the changes in RBP complex composition and subcellular distribution in response to *C9ORF72* amyotrophic lateral sclerosis/frontotemporal dementia dipeptide repeats and DNA damage stress. The coRIC map provides a valuable new approach for defining the roles of RBPs in homeostasis and disease.

## Main Text

Biomolecular interactions, encompassing those between DNA, RNA, and proteins, are fundamental for biology. Despite their significance, RNA-protein interactions remain poorly understood compared to other relationships such as DNA-protein or protein-protein. RNA-binding proteins (RBPs), traditionally known as the interconnectors in RNA life cycle regulation, are emerging as wider regulators of cell function^1^. Their distinctive physicochemical properties, particularly the enrichment of intrinsically disordered regions, enable them to form dense networks with other biomolecules in multiple subcellular localizations^1,2^. These networks are fine-tuned by internal and environmental signals that modulate the interaction affinities between components, for example through post-translational modifications, conferring plasticity. These characteristics enable broad functionality that may include bridging cellular processes spanning different compartments, moonlighting roles in specific locations and regulation of cellular architecture. For example, the exon-junction complex components in the nuclear speckle mediate co-transcriptional splicing but when transported to the cytoplasm bound to spliced mRNAs they cooperate with other partners to regulate nonsense-mediated mRNA decay during translation^3^. As another example of an important additional function, phosphorylation of the RNA processing factor TDP-43 promotes cytoplasmic transportation to induce stress granule formation, a type of phase-separated membraneless organelle that sequesters and stores ribonucleoprotein (RNP) complexes for use in multiple processes^4^. Notably, TDP-43 mutations result in the formation of insoluble cytoplasmic aggregates that hijack many key cellular regulators in amyotrophic lateral sclerosis/frontotemporal dementia (ALS/FTD) patients.

Systematic elucidation of the subcellular localization of RBPs and their biomolecular networks is essential to understanding their function in homeostasis and disease. In recent years, a series of strategies have been developed to interrogate the subcellular localization of RBPs. These are generally based on either fractionation^5,6^ or inference from comparisons of RNA interactome datasets with subcellular proteome atlases^7–9^. Fractionation has a poor signal-to-noise ratio and can only be applied to selected cellular compartments, whereas inference does not provide direct measurement of location. Moreover, both these approaches provide a static snapshot and cannot be easily deployed to study temporal aspects of RBP intercompartmental crosstalk. Thus, new techniques and analytical pipelines are needed to move the field forward. Here, we describe the coRIC map, a multimodal experimental resource and analytical pipeline for studying subcellular RBPs, both in the steady state and under perturbation. The coRIC map datasets are generated using an optimized combination of proximity labeling and phase-separated RNA interactome capture. RBPs are then incorporated into a multidimensional space and integrated with protein-protein interaction (PPI) information to determine the hierarchy of RBP-containing complexes across the cell at multiple scales.

## Results

### Generating a subcellular RBP map

We set out to generate a subcellular RBP map that can be used to study RBP dynamics at the whole cell level. We envisaged that isolating RBPs with subcellular resolution, high sensitivity and dynamicity would require: 1) an optimal protocol combining the efficient pull-down of RNP complexes with proximity labeling of these biomolecules using engineered protein baits associated with known cell locations, 2) integration of all RBP preys into a continuous multidimensional space.

To achieve this, we combined orthogonal phase separation RNA interactome capture (OOPS)^10^, which can pull down RBPs bound to any RNA species, with apurinic/apyrimidinic endonuclease 2 (APEX2)-based proximity labeling^11^. APEX2 was chosen because it allows robust biotinylation in a relatively short time. Zero-distance crosslinking between individual RBPs and RNA was induced by UV exposure^12,13^. Multiple parameters were systematically optimized to ensure efficient RBP pulldown with minimal background (see Materials and Methods; **Fig. 1a and Extended Data Figs. 1 and 2**). We termed this modified approach *<u>co</u>mpartmentalized <u>R</u>NA Interactome <u>C</u>apture* (coRIC) and applied it to the HEK293T cells using 22 baits that cover multiple cellular locations (**Fig. 1b**). This cell line was selected as it is widely used in biomedical research and extensive reference datasets are available for comparison and integration. For the endoplasmic reticulum (ER) lumen, we use horseradish peroxidase (HRP) because it has higher specificity than APEX2 for this compartment^14^. Immunofluorescence for each bait cell line was performed to confirm the expected localization and APEX activity (**Extended Data Fig. 3**). The reproducibility of coRIC samples using each bait was high across replicates (**Extended Data Fig. 4**).

**Fig. 1.**
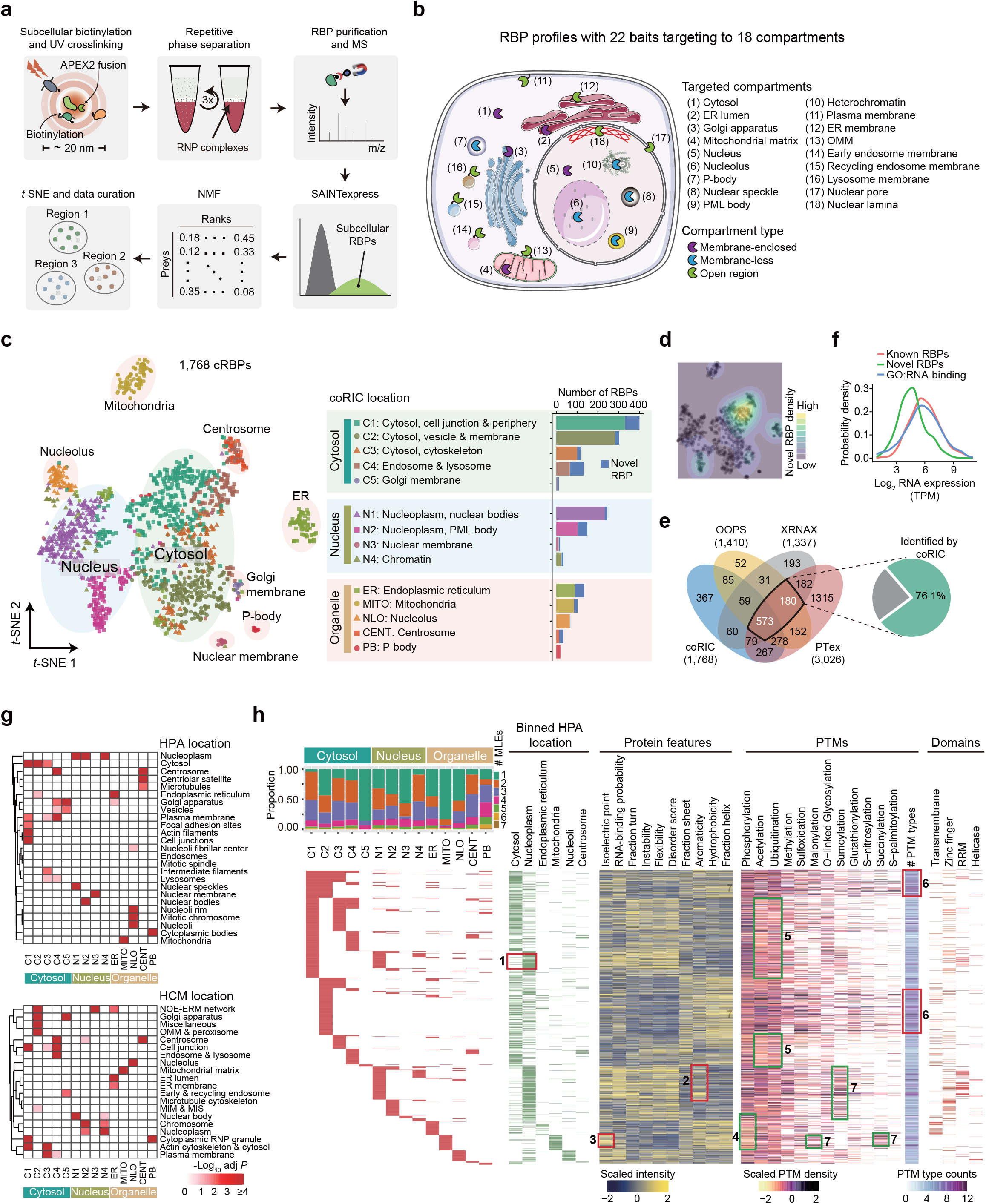
Generation of a compartmentalized RBP map. **a**, Schematic showing the coRIC methodology and analytical pipeline. RNP, ribonucleoprotein; MS, mass spectrometry. NMF, non-negative matrix factorization. *t*-SNE, *t*-distributed stochastic neighbor embedding. **b,** Overview of the targeted compartments for RBP profiling by coRIC. Each bait was N- or C- terminally fused to the APEX2 and FLAG/V5/GFP tag. ER, endoplasmic reticulum; OMM, outer mitochondrial membrane. **c,** *t*-SNE plot showing the subcellular distribution of cRBPs (left). The coRIC locations were named based on each compartment’s representative GO cellular component term/terms. Bar plot showing the number of known and novel RBPs in each compartment (right). cRBPs, coRIC-identified RBPs. **d,** Density plot showing the distribution of the novel RBPs identified by coRIC on the *t*-SNE plot from (c). **e,** Venn diagram comparing the 1,768 cRBPs with different phase separation RNA interactome capture methods (left), the black-outlined box highlights the number of RBPs shared across phase separation–based approaches. The pie chart shows the percentage of shared RBPs identified by coRIC (right). OOPS, orthogonal organic phase separation; XRNAX, protein-xlinked RNA extraction; PTex, phenol toluol extraction. **f,** Density histogram showing the average mRNA expression levels of novel RBPs and known RBPs identified by coRIC and proteins with ‘RNA binding’ annotation in the GO database. TPM, transcripts per million. **g,** Heatmap showing the enrichment of cRBPs detected by coRIC in the corresponding location of HPA^20^ (upper) and HCM^21^ (bottom). The color intensity represents −log10 adjusted *P* value and indicates the degree of enrichment (Fisher’s exact test). HPA, human protein atlas; HCM, human cell map; NOE-ERM network, nuclear outer membrane-ER membrane network; MIM & MIS, mitochondrial inner membrane & mitochondrial intermembrane space. **h,** Heatmap and bar plot summarizing the cRBP characteristics. Bar plot (upper-left) showing the proportion of MLEs for cRBPs in each compartment. Heatmap showing the coRIC- or binned HPA-assigned localization (left), protein features (center-left), PTM density and number of PTM types (center-right), and selected protein domains (right) for all cRBPs. Box 1, dual nucleus-cytosol cRBPs in coRIC displaying exclusive nuclear localization in HPA. Box 2, nucleus single-localized cRBPs have higher levels of aromatic residues. Box 3, mitochondrial cRBPs have higher isoelectric point. Box 4, cRBPs from mitochondria and ER have lower phosphorylation density. Box 5, cytoplasmic cRBPs except for the vesicles & membrane have lower ubiquitination and acetylation density. Box 6, vesicles & membrane cRBPs have multiple types of PTMs. Box 7, nuclear cRBPs have high sumoylation density, whereas mitochondrial cRBPs have higher malonylation and succinylation density. PTM, post-translational modification; RRM, RNA recognition motif.

**Fig. 2.**
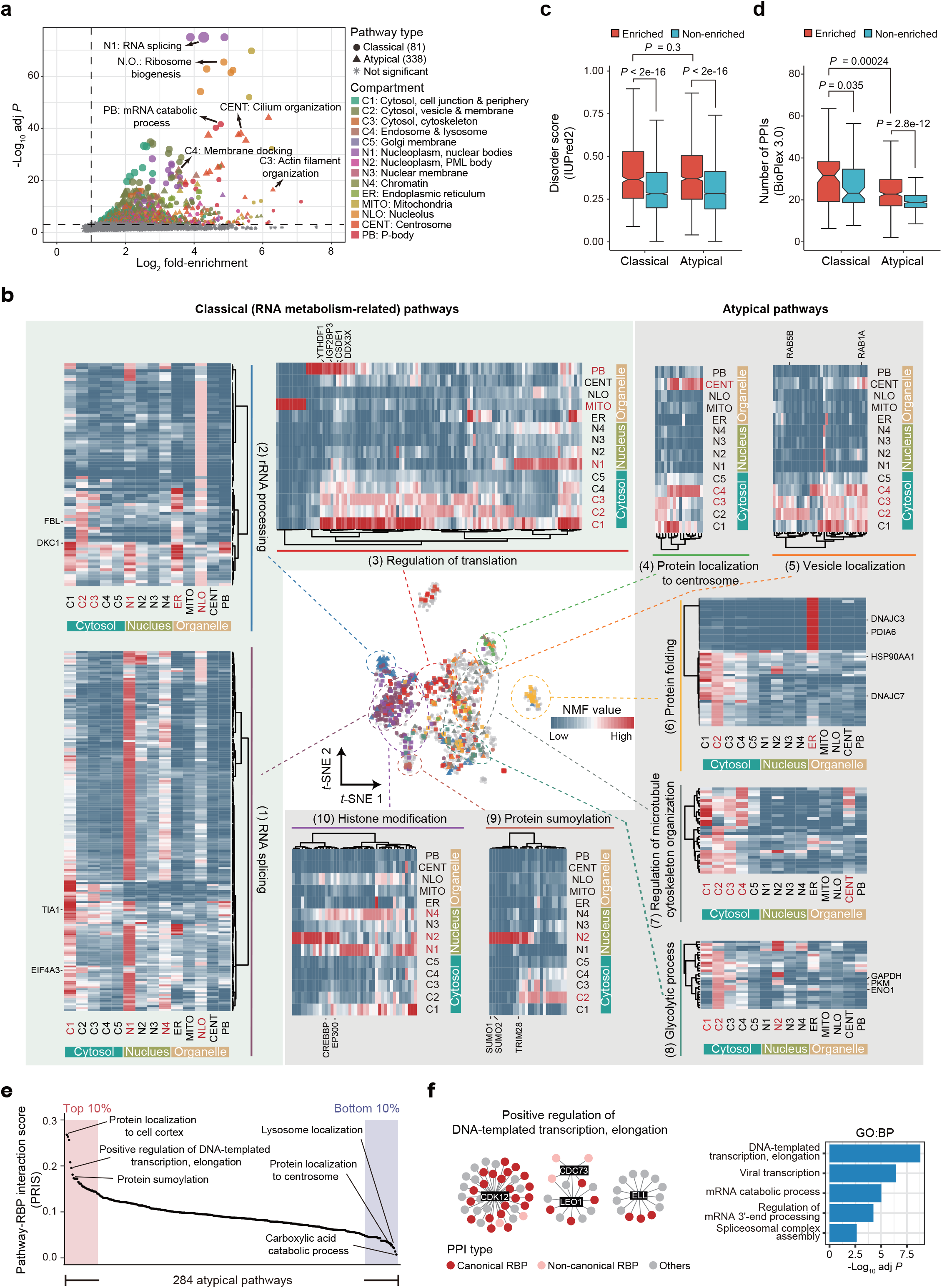
cRBPs participate in multiple classical and atypical biological pathways. **a,** Volcano plot of GO biological process analysis for cRBPs in each compartment. Individual GO terms are represented by either circle or triangle symbols, with their size indicating the number of enriched cRBPs corresponding to each term. The cutoff for enriched terms was set to log2 fold-enrichment ≥ 1 and −log10 adjusted *P* value ≥ 2 (Fisher’s exact test, Benjamini-Hochberg corrected). **b,** Heatmap showing the distribution of cRBPs in selected classical (light green) and atypical pathways (grey). The compartments enriched in the selected pathways are marked in red (Fisher’s exact test, Benjamini-Hochberg corrected *P* ≤ 0.01). The cRBPs participating in the selected pathways are highlighted with colored dots in the *t*-SNE plot (center). **(c-d)** Box plot summarizing the disorder scores (c) and the number of PPIs (d) of the enriched cRBPs and non-enriched proteins in classical or atypical pathways. Disorder scores were predicted using IUPred2. PPI information was taken from BioPlex 3.0 database^51^. *P* values were generated with a two-tailed Mann-Whitney Wilcoxon test. PPI, protein-protein interaction. **e,** Dot plot showing the distribution of the 284 atypical pathways ranked by pathway-RBP interaction score (PRIS) in (Extended Data Fig. 8d). Representative pathways within the top 10% (pink) or bottom 10% (purple) of the score are shown. **f,** PPI networks showing the proteins interacting with the cRBPs in the ‘DNA-templated transcription, elongation’ pathway (left). The black rectangular box indicates pathway-containing cRBPs. The colored circles indicate different types of interactors. Bar plot showing the representative GO biological process terms for the interacting proteins (right) (Fisher’s exact test, Benjamini-Hochberg corrected *P* ≤ 0.01).

**Fig. 3.**
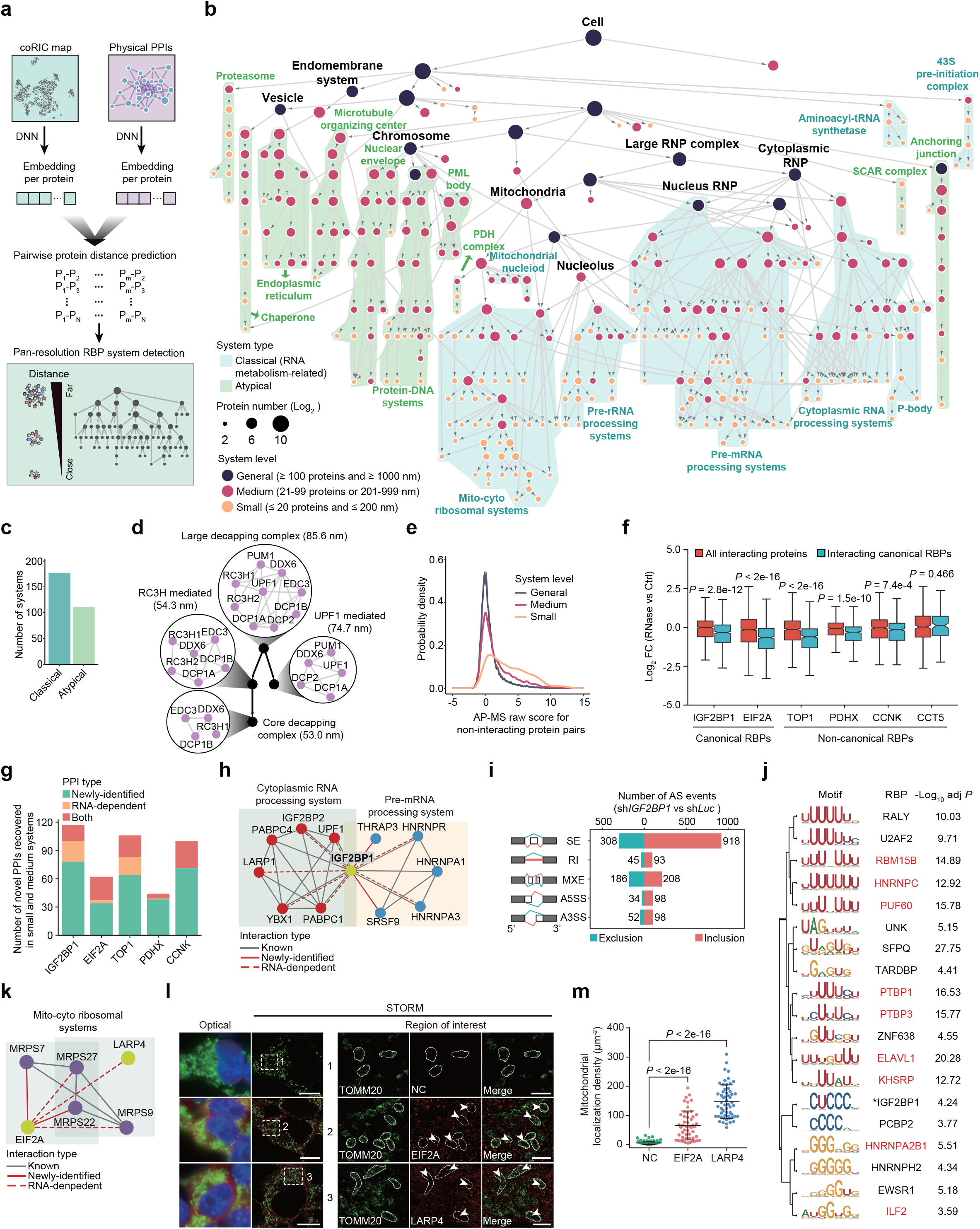
A hierarchical map of subcellular RBP systems across the cell. **a,** Schematic showing the overview of dataset integration strategy. coRIC map and PPI data were analyzed to generate neuronal network embedding for each cRBP. These embeddings were further dissected into an integrated hierarchical network of RBPs inside the cell. DNN, deep neural network. **b,** Visualization of the generated cRBP hierarchical network. Nodes indicate RBP systems, node size represents the number of proteins in the system, node color indicates system levels, and arrow indicates the branches of small systems derived from large systems. The background color defines classical (RNA metabolism-related) and atypical (non-RNA metabolism-related) systems. **c,** Bar plot showing the number of atypical and classical systems in the RBP hierarchical network. **d,** Hierarchical network of decapping complexes in the systems related to P-body. Each circle indicates an RBP system. Nodes and edges inside the circle represent systems-containing proteins and their biophysical interactions, respectively. **e,** Density histogram showing the distribution of AP-MS raw scores from the BioPlex 3.0 dataset for non-interacting protein pairs in different system levels defined in (b). **f,** Box plot showing the abundance of the interactors for each of the selected RBPs with or without RNase treatment. All interactors (red) or interacting canonical RBPs (blue) were grouped for statistic comparison. *P* values were generated with two-tailed Mann-Whitney Wilcoxon tests. Log2 FC, Log2 fold-change. **g,** Bar plot showing the number of novel PPIs in small and medium systems of the RBP hierarchical network that were recovered by AP-MS for each of the selected RBPs. The PPIs were divided into three categories: newly identified (green), RNA-dependent (orange) or both (red). **h,** PPI network of IGF2BP1-containing cytoplasmic RNA processing and pre-mRNA processing system. Nodes represent system proteins and edges represent their biophysical interactions. The biophysical interactions are divided into three categories: known (grey solid line); newly-identified (red solid line), and RNA-dependent (red dashed line). **i,** Bar plot showing the number of alternative splicing (AS) events upon knockdown of IGF2BP1 (|inclusion level difference| ≥ 10 % and FDR ≤ 0.01). Inclusion and exclusion indicate the increasing and decreasing inclusion level of AS events in sh*IGF2BP1* relative to sh*Luc* controls, respectively. SE: skipped exon; RI: retained intron; MXE: mutually exclusive exons; A5SS: alternative 5’ splice site; A3SS: alternative 3’ splice site. **j,** Hierarchical clustering of the RBP motifs associated with the regions of the splice sites belonging to the SE event induced by IGF2BP1 depletion (Fisher’s exact test, Benjamini-Hochberg corrected *P* ≤ 0.001). Only motifs belonging to RBPs that interact with IGF2BP1 are shown. RNA-dependent interactors are highlighted in red. The asterisk (*) indicates IGF2BP1. **k,** PPI network of EIF2A and LARP4-containing mito-cyto ribosomal systems. Nodes represent system proteins and edges their biophysical interactions. The biophysical interactions were divided into 3 categories: known (grey solid line); newly-identified (red solid line), and RNA-dependent (red dashed line). **l,** Left: representative conventional images (optical) and STORM renderings of HEK293T cells stained for LARP4/EIF2A (red), TOMM20 (mitochondrial membrane, green), and DNA (nuclei, blue). Right: zoom of the regions in the white dotted boxes. The white line indicates the outline of the mitochondria. Arrows indicate typical LARP4 or EIF2A foci localized in the mitochondrial matrix. The negative control (NC) sample was stained with no primary antibody. Scale bars: left, 10 μm; right, 2 μm. **m,** Quantification of the mitochondrial matrix localization density of the NC, LARP4, and EIF2A in HEK293T cells with STORM. The density was calculated by dividing the mitochondrial localization signal by the area of the mitochondria (μm^2^). The NC sample was stained with no primary antibody. Data are mean ±standard deviation (s.d.) of n = 125 mitochondria from 23 cells (NC), n = 51 mitochondria from 14 cells (EIF2A), and n = 57 mitochondria from 16 cells (LARP4). *P* values were generated using a two-tailed Mann-Whitney Wilcoxon test.

**Fig. 4.**
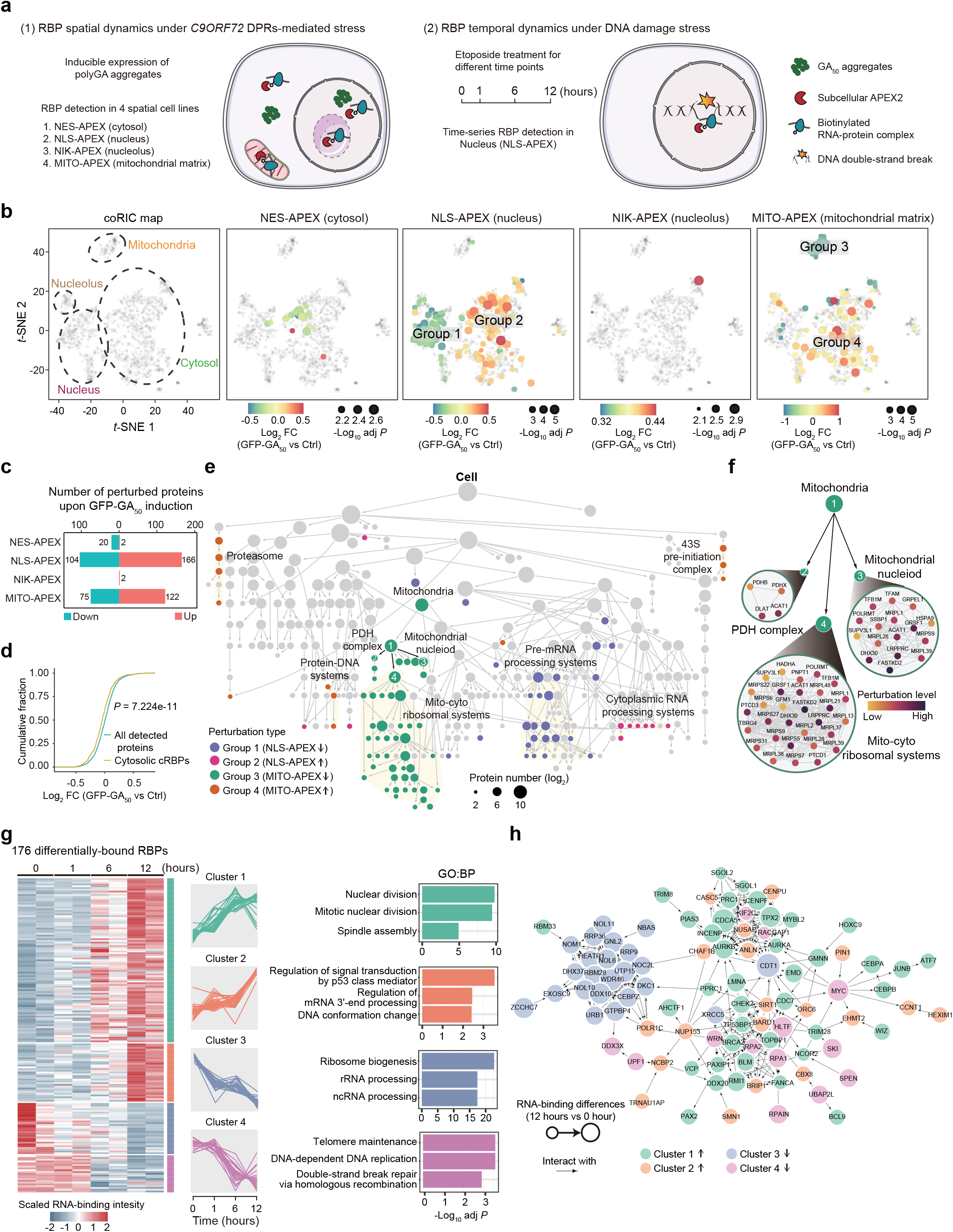
Stress-induced dynamics of cRBPs. **a,** Schematic showing the experimental design to explore RBP spatiotemporal changes when polyGA DPRs are overexpressed (left) or in etoposide-induced DNA damage stress (right) using coRIC. DPRs, dipeptide repeats. **b,** Projection of the significantly changed RBPs induced by polyGA from the indicated APEX-fusion cell lines onto the *t*-SNE map generated in (Fig. 1c). The compartments of interest in the original *t*-SNE plot are marked with dotted ellipses and the changed RBPs in the APEX-fusion cell lines are highlighted with gradient color proportional to the log2 fold-change (moderated *t*-test, Benjamini-Hochberg corrected *P* ≤ 0.01). **c,** Bar plot showing the number of perturbed proteins for each cell line under polyGA induction. Up, upregulated proteins; down, downregulated proteins. **d,** Cumulative distribution of abundance (log2 fold-change) of all detected proteins (blue) and cytosolic cRBPs (yellow) captured by coRIC in NES-APEX cell line upon polyGA induction. *P* value was generated using a Kolmogorov-Smirnov test. **e,** Projection of the significantly changed RBPs caused by the expression of polyGA in the APEX-fusion cell lines onto the RBP hierarchical network in (Fig. 3b). Systems with at least 30% proteins perturbed were colored. **f,** Hierarchical network of the systems labeled in panel E (number 1-4). Nodes and edges represent system-contained proteins and biophysical interactions, respectively. The color of the node indicates the perturbation level, from low to high. **g,** Heatmap of the 176 differentially-bound RBPs induced by etoposide (left) (moderated *t*-test, Benjamini-Hochberg corrected *P* ≤ 0.01). The modulated RBPs were grouped into four clusters based on the scaled RBP abundance using hierarchical clustering. Representative GO biological process terms for each cluster are shown (right) (moderated *t*-test, Benjamini-Hochberg corrected *P* ≤ 0.01). **h,** PPI networks of the 176 differentially-bound RBPs using STRINGDB. The nodes and edges indicate the changed RBPs and STRINGDB-integrated interactions, respectively.

By comparing individual datasets to their respective neighboring spatial reference, we were able to identify RBPs enriched by each bait (**Supplementary Table 1**). This approach evaluated the baits’ performance. For example, 237 RBPs were enriched in MITO-APEX (mitochondrial matrix) compared to NES-APEX (cytosol) cells, with 61% (145/237) recognized as mitochondrial RBPs and 35.4% (84/237) mitochondrial proteins with previously unknown RBP function (**Extended Data Fig. 5a-c**). Western blotting of coRIC lysates confirmed that the mitochondrial RBPs FASTKD2 and GRSF1 and the cytosolic RBPs YTHDF3 and G3BP1 are predominantly captured using the MITO-APEX and NES-APEX cells, respectively (**Extended Data Fig. 5d**). Comparison of nucleus-enriched RBPs isolated by coRIC (NLS-APEX relative to NES-APEX cells) with two reported datasets combining oligo(dT) capture and cellular fractionation (eRIC and seRIC)^5,6^ showed substantial overlap but many RBPs were only present in coRIC (**Extended Data Fig. 5e**). We also compared the nucleolus (NIK-APEX relative to NLS-APEX cells) and outer mitochondrial membrane (OMM-APEX relative to NES-APEX cells) datasets with a recently reported dataset using formaldehyde crosslinking and proximity labeling for subcellular RBP purification^15^. Formaldehyde induces multiprotein covalent bonding and, thus, also captures indirect interactions^16^. Consistently, coRIC showed higher specificity for biological processes and cellular components related to these compartments (**Extended Data Fig. 5f**).

The successful identification of RBPs corresponding to individual compartments encouraged us to integrate all baits into a comprehensive subcellular RBP map, which we term coRIC map. By scoring the protein intensity data of each bait against negative controls using SAINTexpress, we identified 1,768 high-confidence RBPs (hereafter referred to as coRIC-identified RBPs or cRBPs). Next, we extracted the localization information of proteins using non-negative matrix factorization (NMF), dividing these 1,768 cRBPs into 14 subcellular compartments and visualizing this distribution using *t*-distributed stochastic neighboring embedding (*t*-SNE) (**Fig. 1c**). Within each compartment, the NMF value of a specific RBP served as an indicator of its spatial distribution intensity. The compartment displaying the highest NMF value was defined as the primary location of each RBP, whereas values in other compartments provided a relative position compared to the main compartment. The subcellular compartments were defined based on the enrichment of gene ontology (GO) terms associated with the cytosol (C1: cell junction & periphery; C2: vesicles & membrane; C3: cytoskeleton; C4: endosomes & lysosomes; C5: Golgi membrane), the nucleoplasm (N1: nuclear body; N2: PML body; N3: nuclear membrane, N4: chromatin) and the organelles (ER: endoplasmic reticulum; MITO: mitochondria; NLO: nucleolus; CENT: centrosome; and PB: P-body). Projection analysis revealed the distribution of subcellular information derived from the 22 baits in the 14 compartments (**Extended Data Fig. 6a**). The identification of the centrosome as an independent compartment, for which we had not used a specific bait, can be explained by the fact that this organelle is connected to different cellular structures and is included within the cytosol^17^. Consistently, our baits designed for the cytosol, endosome membrane and lysosome membrane captured many centrosome proteins.

Among the 1,768 cRBPs, 19.1% (338) were novel RBPs based on comparison with two integrated RBP datasets (proteins identified in phase separation approaches or with RNA binding annotation in the GO database)^10,18,19^ (**Fig. 1c and Supplementary Table 2**). These novel RBPs were distributed across all 14 compartments but concentrated mainly in the PML body, endosomes & lysosomes, centrosome, and ER (**Fig. 1d**). The identification of a large proportion of RBPs included in phase separation RNA interactome capture datasets^10,18,19^ (**Fig. 1e**), confirmed that coRIC has high sensitivity. Interestingly, we noticed that the 338 novel RBPs display lower mRNA levels compared with RBPs included in other datasets (**Fig. 1f**). This demonstrates that coRIC captures RBPs enriched in specific subcellular locations even when total cellular levels are low, providing a tool for studying RBPs that are missed using conventional approaches. As expected, cRBPs were significantly enriched in the corresponding subcellular regions defined by the Human Protein Atlas (HPA)^20^, the reference imaging dataset for protein subcellular localization, and the Human Cell Map (HCM)^21^, a proximity-dependent biotinylation map of subcellular compartments in HEK293T cells (**Fig. 1g**). For a selected panel of cRBPs, we used photoactivatable-ribonucleoside-enhanced crosslinking and immunoprecipitation (PAR-CLIP) to provide independent validation that they bind RNA (**Extended Data Fig. 6b**).

### Characteristics of cRBPs

To gain insight into both general and specific features of the human RBP repertoire across subcellular locations, we studied the presence of multi-localization events (MLEs), biophysical properties, post-translational modifications (PTMs) and protein domains of the 1,786 cRBPs (**Fig. 1h**, **Extended Data Fig. 7a-e and Supplementary Table 2**).

Proteins often function in multiple subcellular compartments. Closer inspection of the NMF analysis showed that 51% (902) of the cRBPs display MLEs, which is comparable to previous subcellular localization studies: 50% in HPA and 54.3% in hyperLOPIT (hyperplexed localization of organelle proteins by isotope tagging), a high-resolution spatial protein map generated by cellular fractionation^22^. The number of MLEs per protein ranged from 2 to 7 and although they occurred more frequently between nearby locations, we also observed MLEs between distant compartments (**Fig. 1h, left; Extended Data Fig. 7a**). Notably, despite the consistency between the main locations in our coRIC map and the HPA annotations, we observed discrepancies when looking at MLEs. For example, a substantial number of cRBPs (110) with dual nucleus-cytosol signals in coRIC displayed exclusive nuclear localization in HPA. This subset of proteins was enriched for GO terms related to spliceosome and stress granule components, which are known to shuttle between the nucleus and cytoplasm^23^ (**Extended Data Fig. 7b**). Therefore, coRIC can detect MLEs that may be missed in immunofluorescence approaches due to limited dynamic range. We also interrogated the biophysical properties of the cRBPs through analysis of amino acid sequence (**Fig. 1h, center; Extended Data Fig. 7c**). Consistent with the current knowledge of canonical RBPs^24^, cRBPs showed higher hydrophily, disordered scores and predicted RNA-binding probability compared to both total proteome and subcellular proteomes. Yet, we observed differences across subcellular compartments. Nuclear RBPs had higher disordered scores and predicted RNA-binding probability compared with cytosolic RBPs, which may contribute to the formation of larger RBP-mediated protein complexes in this compartment^25^. Nucleus single-localized RBPs also had higher levels of aromatic residues than nucleus-cytoplasm dual-localized RBPs, which is crucial for enhancing the stability of protein-protein interactions^26^. Within the cytoplasm, mitochondrial RBPs had higher isoelectric point compared to other compartments, which may represent an adaptation to the alkaline environment of the mitochondrial matrix^27^.

Next, we studied the relative frequency of PTMs among cRBPs by matching our map to the dbPTM database^28^ (**Fig. 1h, center-right; Extended Data Fig. 7d**). Most classical PTMs were more abundant in cRBPs compared to both total proteome and individual subcellular proteomes, with some modifications exhibiting a preference for proteins in specific subcellular locations. Phosphorylation was prevalent across all compartments except for the mitochondria and ER. Acetylation and ubiquitination were widely distributed but displayed lower levels in the cytoplasmic compartments except for the vesicles & membrane. In general, RBPs in the vesicle & membrane tended to have multiple types of PTMs, which may be associated with their dynamic role in cellular trafficking and communication. Sumoylation was more enriched in the nucleoplasm, while succinylation and malonylation were more enriched in the mitochondria.

Finally, we investigated the abundance of protein domains to shed light on the enrichment of RNA-binding domains (RBDs), specific types of proteins and functions (**Fig. 1h, right; Extended Data Fig. 7e-h and Supplementary Table 4**). Nuclear RBPs contained a higher fraction of well-characterized RBDs (RNA recognition motif [RRM] and zinc finger). Moreover, we observed many non-canonical RBDs enriched in certain compartments, such as thioredoxin and PDZ domains in the ER and cell junction & periphery, respectively^29^. Interestingly, we also noticed transmembrane domains such as the LEM domain in the nuclear membrane and small GTPase domain in the Golgi membrane, among others. This was intriguing because although integral membrane proteins (memRBPs) have been recognized as RBPs^30^ they remain largely unstudied. We reasoned that the combination of OOPS and proximity-labelling enabled the capture of relatively insoluble memRBPs that are otherwise difficult to isolate using conventional RNA interactome methods. Supporting this idea, we projected our dataset onto the hyperLOPIT fractionation map and found that many RBPs were enriched in membrane-associated regions (**Extended Data Fig. 7f**). The number of memRBPs detected by coRIC was larger than in other RNA interactome studies (**Extended Data Fig. 7g**). These memRBPs were enriched in functions related to the GTPase cycle and protein processing in the ER, as shown using Metascape (**Extended Data Fig. 7h**). We validated two of them, CLCC1 and POR, as RBPs using PAR-CLIP (see above **Extended Data Fig. 6b**).

In summary, we have developed a new methodology and an analytical pipeline to study RBPs that provides spatial resolution in an integrative and dynamic manner. This approach offered insights into the distinctive characteristics of RBPs in subcellular locations.

### Understanding the functional diversity of cRBPs in biological pathways

To interrogate the putative biological roles of the 1,768 identified cRBPs in their respective subcellular compartments, we used GO biological process analysis, including the MLEs for each respective cRBP. This yielded 419 significantly enriched biological pathways that could be divided into 81 RNA metabolism-related (hereafter termed classical; 725 RBPs) and 338 non-RNA metabolism-related (hereafter termed atypical; 1,043 RBPs) pathways, distributed across all targeted cellular compartments, with the top 100 categories more enriched in classical pathways (66 GO terms) (**Fig. 2a**, **Extended Data Fig. 8a and Supplementary Table 3**). Individual atypical pathways generally contained a smaller fraction of cRBPs than classical pathways (**Extended Data Fig. 8b**). As expected, classical pathways included GO terms such as ‘RNA splicing’ in the nucleus, ‘ribosome biogenesis’ in the nucleolus, and ‘mRNA catabolic process’ in the P-body. Atypical pathways were more abundantly distributed in cytoplasmic regions and included for example ‘cilium organization’ in the centrosome, ‘membrane docking’ in the endosome and lysosome, and ‘actin filament organization’ in the cytoskeleton, highlighting the wide functional diversity of the identified cRBPs (**Extended Data Fig. 8c**). Our RBP-centric cellular pathway analysis offers a valuable opportunity to interrogate how classical and atypical pathways and their constituent RBPs involve unexpected cellular locations. This is relevant to understand how pathway execution involves collaboration between subcellular compartments^31^. It can also help identify potential moonlighting functions for specific RBPs outside their main subcellular compartment.

We explored the distribution of cRBPs involved in classical pathways (**Fig. 2b, left**). As examples, nucleolus rRNA modification enzymes (DCK1 and FBL) were localized in the ER, cytosolic translation regulators (DDX3X and CSDE1) in the P-body, and nuclear splicing factors (TIA1 and EIF4A3) in the cytoplasm. Although splicing factors are predominantly nuclear, they frequently translocate to the cytoplasm to regulate other aspects of RNA metabolism^23^. Similarly, rRNA processing can occur in other compartments apart from the nucleolus, such as m3U methylation in the cytoplasm^32^. Likewise, protein translation regulators have been co-purified with either P-bodies or core P-body structural proteins^33,34^, which may indicate a role in re-establishing translation after mRNAs exit P-bodies.

In the atypical pathways **(Fig. 2b, right)**, we identified previously reported moonlighting RBPs localized to multiple compartments. As examples: we detected CREB-binding protein (CREBBP) in the nucleus, where it binds to enhancer RNAs to stimulate histone acetyltransferase activity^35^; the cytosolic metabolic enzyme pyruvate kinase muscle (PKM), which exerts a non-canonical function as a protein translation regulator by interacting with subsets of ER-associated ribosomes^36^; the E3 SUMO-protein ligase tripartite motif containing 28 (TRIM28) recruited by transposable element-containing *LINE-1* RNAs to deposit the repressive H3K9me3 histone mark in mouse embryonic stem cells^37^; and protein disulfide isomerase 6 (PDIA6), which regulates cancer progression through unclear RNA-related mechanisms^38^. We also identified RBPs involved in vesicle trafficking and cytoskeleton organization, in line with knowledge that cytoplasmic vesicles regulate RNA intracellular transport^39^. Among them, we observed RAB5, which is part of the RAB5 effector FERRY complex that links mRNAs to early endosomes for axonal transport^40^, and RAB1, which is involved in internal ribosome entry site (IRES)-mediated RNA localization to the ER for translation^41^. Apart from these reported cases, we identified many core centrosomal subunits as novel RBPs and validated three of them (ODF2, CEP135 and POC5) using PAR-CLIP (see above **Extended Data Fig. 6b**). Interestingly, although the centrosome associates with multiple RBPs in neurons^42^, it is also well-established that the centrosome structure is disrupted by RNase treatment^43^. We speculate that the interaction of core centrosomal subunits with RNA regulates centrosome assembly and/or maturation in an RNA-dependent manner.

Given the substantial involvement of cRBPs in atypical pathways, we investigated whether these proteins possess unique features that support their function. The overall disorder score of the cRBPs in atypical pathways was comparable to RBPs in classical pathways but substantially higher than non-enriched proteins in the same pathways (**Fig. 2c**). This suggests that these cRBPs have the potential to engage in multivalent interactions similar with RBPs in classical pathways^44^. Supporting this idea, cRBPs in both classical and atypical pathways were involved in a larger number of PPIs than non-enriched proteins in the same pathways (**Fig. 2d**). We then designed a ranking score for calculating the degree of the RBP interactions for cRBPs in each atypical pathway, which we named pathway-RBP interaction score (PRIS) (**Extended Data Fig. 8d**). Among the top 10% PRIS-ranked pathways, we identified for example ‘positive regulation of DNA-templated transcription, elongation’, ‘protein localization to cell cortex’ and ‘protein sumoylation’, and among the bottom 10% ‘lysosome localization’, ‘carboxylic acid metabolic process’ and ‘protein localization to centrosome’ (**Fig. 2e,f and Extended Data Fig. 8e,f**). PPI networks in the top 10% contained more canonical RBPs. Accordingly, the GO biological process terms for the interacting proteins in atypical pathways showed enrichment in functions related to RNA metabolism (regulation of mRNA processing and RNA splicing). In this regard, the core elongation factor CDK12 in ‘positive regulation of DNA-templated transcription, elongation’ has been reported to promote breast cancer cell invasion through its interaction with the splicing machinery^45^. As for ‘protein localization to cell cortex’, the presence of splicing factors interacting with scaffolding factors such as EPB41L2^46^ suggests a role in the transportation of RNA regulatory factors along neuronal axons and dendrites, which is a poorly understood process. In contrast, pathways in the bottom 10% such as ‘lysosome localization’ clustered more with non-canonical RBPs and other proteins related to the same pathway including autophagy and mTOR signaling. These instances illuminate two potential differential modes of action for cRBPs within atypical pathways. First, cRBPs may act as linkers between the corresponding atypical pathway and RNA metabolism through interaction with canonical RBPs. Second, cRBPs may act as scaffolds for other proteins in the same or related atypical pathways, potentially stabilizing their function. In both scenarios, RNA could play an important role.

Our coRIC map provides a template for investigating the biological pathways associated with RBPs in different subcellular compartments and will facilitate a deeper understanding of RBP network organization and function within the cell.

### Dissecting the hierarchy of RBP complexes across the cell

Although GO and PPI datasets provide valuable information about RBP networks, they do not reflect the hierarchy of RBP complexes within those networks. The latter is fundamental to fully understand both individual RBP functions and the crosstalk between RBPs in different complexes^47^. Recent advances in machine learning-based approaches such as MuSIC and NeST^48–50^ dissect the composition of protein complexes across the cell by integrating multimodal complementary datasets with remarkable precision. We envisaged that applying a similar approach to integrate our coRIC map with a large-scale PPI dataset such as BioPlex^51^ could dissect the hierarchies of RBP-containing complexes. Both coRIC and BioPlex were generated from HEK293T cells and share 1,465 RBPs with subcellular distribution and biophysical association information. We used the neuronal network method node2vec to embed each RBP from the two datasets into a unified dimension vector that estimated the pairwise protein distances by random forest regression (**Fig. 3a**). This allowed the systematic identification of RBP communities at multiple scales from small protein complexes to large cellular pathways by progressively relaxing the distance (range from 7 nm to 9 μm). The sensitivity of community detection was further adjusted by comparison with the HCM and the collection of mammalian protein complexes in CORUM^52^ (**Extended Data Fig. 9a**). The optimal hierarchy was then determined by measuring the maximal known protein-pairs within the minimal number of communities (systems), which yielded 308 RBP systems organized into 420 cascading relationships (**Fig. 3b and Supplementary Table 4).** We further divided the 308 RBP systems into large, medium, and small systems based on both the constituent protein numbers and the estimated complex sizes; small systems represented protein pairs with the strongest relationship. These systems comprised intermolecular relationships within RNA metabolism-related (hereafter termed classical; P-body, pre-mRNA processing complex and 43S pre-initiation complex) and non-RNA metabolism-related (hereafter termed atypical; chaperone, microtubule organizing center and PML body) biological processes. The number of classical systems was larger than the atypical and they displayed more branching and frequent interconnections (**Fig. 3c**). This observation is consistent with the fact that RBPs often interact with each other^53^ and participate in multiple steps of the RNA life cycle^54^. As previously reported^48^, systems belonging to the cytoplasmic and mitochondrial ribosome machinery clustered together (mito-cyto ribosomal systems), supporting the abundant crosstalk between both processes.

Our integrative analysis provides a hierarchical network view of RBP complexes across the cell. For example, the mRNA decapping proteins DCP1B, DDX6, EDC3 and RC3H1 were associated under an estimated distance of 53 nm (**Fig. 3d**). As the distance was relaxed (54.3 nm), this complex expanded into a larger one including DCP1A and RC3H2, consolidating with a distinct community that contains the core non-sense decay factor UPF1, the deadenylation mediator PUM1 and other related regulators to form a large (85.6 nm) mRNA decapping complex involved in controlling mRNA decay. We also identified multifaceted RBPs involved in RNA metabolism-related complexes (**Extended Data Fig. 9b**), such as the deubiquitylase USP42 in nuclear speckle-related complexes, or the E3 ubiquitin ligases RNF219 and PRRC2B in CCR4- NOT and 43S pre-initiation complexes, respectively, consistent with supporting evidence in recent reports^55–57^. Likewise, the chromatin remodeling proteins SMARCA4, SS18, ARID1A and ARID1B complexed together, which might be related to the role of non-coding RNAs in maintaining the structure of SWI/SNF complexes^58^. The concordance with independent studies confirms the reliability of our RBP hierarchical network.

In addition, we identified many previously unnoticed components of protein complexes in both classical and atypical systems, with overrepresentation in the mito-cyto ribosomal and pre-mRNA processing systems (**Extended Data Fig. 9c and Supplementary Table 4)**. In this regard, AP-MS (affinity purification-mass spectrometry) raw scores for protein pairs not classified as interacting in the BioPlex dataset were substantially higher in the small systems of our RBP hierarchy than in medium and large ones (**Fig. 3e**). This suggests that our approach may detect certain PPIs that were not defined in BioPlex. We then chose six RBPs involved in different systems for validation, including two classical (EIF2A and IGF2BP1) and four atypical (TOP1, PDHX, CCNK and CCT5) RBPs. Among them, EIF2A, IGF2BP1, TOP1 and CCNK were not previously described as part of their respective RBP complexes. We performed AP-MS for all six RBPs in the presence or absence of RNAse to study the role of RNA in maintaining the structure of their respective complexes^59^ (**Fig. 3f and Supplementary Table 5**). Except for CCT5, all other RBPs had substantial RNA-dependent reductions in their interaction signals. Many of the newly-identified and/or RNA-dependent interactors were present in the corresponding complexes of our RBP hierarchy (**Fig. 3g**). Among the five RBPs with substantial RNA-dependent interactions, we found the involvement of IGF2BP1 in pre-mRNA processing particularly intriguing (**Extended Data Fig. 9c)**, because, despite being known to shuttle between nucleus and cytoplasm^60^, IGF2BP1’s nuclear function is poorly understood. Further exploration of the AP-MS data confirmed that IGFBP1 not only interacts with cytosolic RNP complexes but also with several splicing factors to form a separate mRNA splicing complex (**Fig. 3h**), and interactions with most of these splicing factors were RNA-dependent. Consistent with an important function in splicing, knockdown of *IGF2BP1* induced 2,040 alternative splicing events, and among the types of splicing change, exon skipping events substantially increased (**Fig. 3i**, **Extended Data Fig. 9d and Supplementary Table 6**). This was validated by semi-quantitative PCR for *DMPK* and *SEC31B*, which showed higher levels of exon inclusion in *IGF2BP1*-deficient cells relative to the control (**Extended Data Fig. 9e,f**). The regions contained in these exon skipping events were enriched in RBP motifs for either IGF2BP1 itself or the interacting partners in our AP-MS (**Fig. 3j**), further supporting a role in regulating splicing.

We also studied EIF2A and LARP4, as, unexpectedly, our RBP hierarchy predicted that both cytosolic translation regulators cluster together with mitochondrial ribosomal small subunits as part of the mito-cyto ribosomal systems (see above **Extended Data Fig. 9c**). Similarly, AP-MS confirmed that EIF2A interacts with multiple mitochondrial ribosomal small subunits (MRPS7/9/22/27) in both an RNA-dependent and independent manner (**Fig. 3k**). EIF2A was also localized to mitochondria in the HPA, but the exact position within the mitochondria was unclear, whereas LARP4 is known to be located on the outer mitochondrial membrane^61^. Because MRPS subunits are almost exclusively inside the mitochondrial matrix to assist with mitochondrial translation, we performed STochastic Optical Reconstruction Microscopy (STORM)^62^ in HEK293T and HeLa cells to study in greater detail the distribution of EIF2A and LARP4. As expected, the main signals were in the cytosol, but we also detected specific signal inside the mitochondrial matrix enclosed by the mitochondrial membrane marker TOMM20 (**Fig. 3l,m and Extended Data Fig. 9g,h**). This confirms the physical presence of LARP4 and EIF4A inside the mitochondrial matrix and suggests a relationship with mitochondrial translation.

Our multiscale RBP hierarchy thus provides a valuable resource to generate new hypotheses concerning multifaceted RBP function and can be used to study the role of RNA in maintaining the corresponding biomolecule networks.

### Dynamic reorganization of cRBPs under stress

Studying the remodeling of subcellular RNA-protein interactions is essential for understanding their role in homeostasis and disease. Although recent reports have profiled RBPs during viral infection^63^, oxidative stress^18,64^, cell cycle arrest^10^ and others, these studies primarily focus on whole cell lysates and provide little or no subcellular information. To showcase coRIC’s utility for studying the dynamic rearrangements of RBPs in the subcellular space under cell stress conditions, we investigated the consequences of two disease-relevant treatments (**Fig. 4a and Supplementary Table 7**): 1) *C9ORF72* amyotrophic lateral sclerosis (ALS)/frontotemporal dementia (FTD) dipeptide repeats (DPRs) and 2) DNA double-strand breaks (DSBs).

Alterations of RNA metabolism play a fundamental role in ALS/FTD^65^. DPRs resulting from translation of the *C9ORF72* G4C2 repeat expansion are considered important effectors of neuronal toxicity in these patients but their effects on RNA metabolism are as yet poorly understood^66^. We chose poly glycine-alanine (GA, polyGA) because it is the most abundantly detected DPRs in ALS/FTD patient tissue^67^. To explore whether polyGA DPRs induce changes in RBP subcellular localization, we over-expressed synthetic constructs containing 50 GA dinucleotide repeats fused to the C-terminus of GFP (GFP-GA50)^68^ in an inducible manner in NES-APEX (cytosol), MITO-APEX (mitochondrial matrix), NLS-APEX (nucleus) and NIK-APEX (nucleolus) HEK293T cell lines (**Fig. 4a, left**). Transient expression of polyGA for 24 hours resulted in bright cytoplasmic inclusions without noticeable effect on cell proliferation (**Extended Data Fig. 10a,b**). This enables the capture of early changes in RBP reorganization and function before severe cell phenotype changes occur. We then performed coRIC and determined the total proteome in these cell lines under these conditions using tandem mass tag (TMT) quantification. Consistent with the lack of effect on cell proliferation, the total proteome was only slightly changed by treatment with polyGA (**Extended Data Fig. 10c**).

To explore the spatial changes in RBP dynamics caused by polyGA induction, we compared the differentially captured cRBPs from untreated and polyGA-treated cell lines. This yielded 491 RBPs, which were projected onto our coRIC map. Unexpectedly, although polyGA aggregates are predominantly cytoplasmic, most of the RBP modulation occurred in the NLS-APEX (270) and MITO-APEX cell lines (197), while NES-APEX (22) and NIK-APEX (2) cell line showed fewer overall changes (**Fig. 4b,c)**. Both NLS-APEX and MITO-APEX cell lines captured more RBPs annotated as cytosolic in untreated cells, accompanied by a decrease in those from their respective target compartments. Thus, polyGA causes redistribution of RBPs into alternate compartments. Among the RBPs decreased in NLS-APEX cells we identified HNRNPA3, which is known to be mis-localized in ALS/FTD patient tissue^69^. The most substantially reduced cytosolic RBPs in NES-APEX cells included proteins involved in stress granule formation and translation regulation (YTHDF1 and EIF4B) (**Extended Data Fig. 10d**). However, the number of reduced RBPs in NES-APEX cells was less than anticipated. We reasoned that this is due to the high basal levels of these proteins in the cytosol, which makes changes in this compartment less noticeable. In agreement, there was a trend towards a global reduction of cytosolic RBPs after polyGA induction compared to the control (**Fig. 4d**).

To clarify the RNA regulatory functions potentially affected in NLS-APEX and MITO-APEX cells due to the polyGA-induced RBP redistribution, we divided the changed RBPs into four different groups (two per cell line) and projected them onto our RBP hierarchical network to understand what complexes and functions are disturbed (**Fig. 4e,f**). Group 1 and 2 represented changed nuclear and cytosolic RBPs in NLS-APEX cells, respectively. The former was enriched in molecular complexes containing pre-mRNA processing (SRSF1/2 and HNRNPAB) and the latter in cytosolic translation factors (EIF4G and PABPC4). Group 3 and 4 represented altered mitochondrial and cytosolic RBPs in MITO-APEX cells, respectively. Group 3 was enriched in proteins involved in mitochondrial gene expression and organization. For instance, we observed that within the mitochondria-related hierarchy system, polyGA disrupted systems encompassing the mitochondrial nucleoid complex involved in transcriptional regulation, the PDH complex involved in carbon metabolism, and the mitochondrial translational machinery in the mito-cyto ribosomal system. Group 4 was enriched in cytosolic translation factors (EIF3E and EIF4F) and proteasome regulators (PSMC3 and GOPC). Overall, these findings correlate with recent reports describing a disrupting effect of polyGA on nucleocytoplasmic transport^70^, mitochondria function^71^ and protein homeostasis^72^. We postulate that mis-localization of cytosolic RBPs inside various subcellular compartments induced by polyGA aggregates perturb native RNA-protein interaction networks through direct competition for factors, ultimately leading to abnormal cell function and toxicity.

RBPs act as important effectors of the DNA damage response by guiding the formation of a complex network of biomolecules including DNA, RNAs and proteins to DNA damage foci^73^. How this is regulated is poorly understood. We performed coRIC and total proteome analysis using TMT quantification in NLS-APEX cells treated with the DNA topoisomerase inhibitor etoposide to induce DNA double-strand breaks (DSBs) at a series of time points (0, 1, 6 and 12 hours) (**Fig. 4a, right; Extended Data Fig. 11a**). From the proteome data, we identified a total of 1,087 differentially expressed proteins at any time point compared to the control (**Extended Data Fig. 11b**). As expected, among them there was an increase in DNA damage repair and cell cycle regulators, while cell growth and rRNA processing factors were decreased. To define the differentially modulated RBPs, we selected those that changed at any time point and were present either in the nuclear compartment of our coRIC map or identified as nuclear proteins in the HPA. This yielded 176 differentially bound RBPs, of which 139 (79%) were contained in the DNA damage-NET database (a curated database of DNA damage-related proteins)^74^ (**Extended Data Fig. 11c**), supporting the validity of our strategy. Moreover, 115 (65.3%) were unchanged in the total proteome **(Extended Data Fig. 11d)**, demonstrating that DNA damage alters the ability of these proteins to interact with RNA rather than modifying their expression.

To further understand RBP complex dynamics, we analyzed the kinetics of RBP modulation following etoposide treatment, observing four main clusters (**Fig. 4g,h**). Clusters 1 and 2 were larger (126 proteins) and contained RBPs increasing at 6 and 12 hours, respectively. Among them, 70 were novel RBPs (**Extended Data Fig. 11e**), indicating *de novo* RNA-protein interactions induced by DNA damage. These proteins were enriched in GO biological function terms related to chromatin regulation such as the nuclear division, DNA conformation change, and p53 signaling. This is interesting because dual DNA-RNA binding proteins (DRBPs) are considered important regulators of the DNA damage response^75^. For example, we observed increasing RNA-binding of two DNA repair factors TP53BP1 and BRCA2, both of which are recruited by RNA to DNA damage foci for safeguarding efficient double-strand repair^76,77^. Cluster 3 represented RBPs reduced at one hour and was enriched in ribosome biogenesis factors, most of which were not decreased at the protein level (**Extended Data Fig. 11f**). A recent study described the recruitment of pre-rRNAs from the nucleolus to DNA damage foci to promote DNA repair^78^. We asked whether our-identified ribosome biogenesis factors are recruited to DNA damage loci or instead stay unbound to RNAs in the nucleoplasm. Laser microirradiation assay^79^ demonstrated that all five selected candidates (DDX10, GNL2, NOL10, GTPBP4 and RRP36) accumulate at DNA damage loci upon irradiation and rapidly dissociate (**Extended Data Fig. 11g**). This suggests that the stoichiometry of the interaction between ribosome biogenesis factors and RNAs at DNA damage foci is reduced during DNA double-strand break stress compared to the nucleus in basic conditions. These ribosome biogenesis factors may recruit pre-rRNAs to DNA damage foci and/or participate in DNA repair, which would need to be further explored. Cluster 4 corresponded to RBPs showing a more progressive reduction and contained multiple RNA-DNA hybrid regulators including the DNA repair and damage checkpoint activation factor RPA1 and RPA2 and the core nonsense-mediated decay factor and RNA/DNA helicase UPF1, all with decreased RNA-binding not caused by reduced protein levels (**Extended Data Fig. 11h**). The decreased interaction of these factors with RNA might be either a consequence or a facilitator of the cleavage of R-loops at DNA damage foci^80,81^.

These results demonstrate that the coRIC map can be applied to study the spatiotemporal dynamics of RBPs under stress, which can help unveil disease-relevant mechanisms.

## Discussion

The traditional view of eukaryotic cellular processes occurring in a reaction-diffusion manner inside relatively large compartments, has been debunked by the realization of high levels of modular organization within specific subcellular domains^82^. RBPs are a multifaceted part of the proteome that plays pivotal roles in cellular compartmentalization through multivalent interactions. Yet, our comprehension of the underlying phenomena remains limited. We have generated the coRIC map as a resource and analytical framework for charting subcellular RBP characteristics, and their participation in pathways and biomolecular networks in a spatially resolved manner. Our cRBP purification approach is based on the optimized combination of APEX-mediated proximity labeling^11^ and phase-separation RNA interactome capture^10^ and has high resolution and sensitivity. Visualization of coRIC datasets within a continuous virtual space allowed us to identify relative distances between RBPs in different locations to create a RBP map. This represents a major advancement over other methodologies and pipelines that do not provide direct spatial information. Using this approach, we have identified 1,768 RBPs in 14 subcellular compartments. These cRBPs display both common and specific features depending on their respective localizations and show highly interactive intercompartmental crosstalk whose deeper understanding will help shed light on their function.

Our coRIC map provided multiple interesting observations. For example, we identified a large group of memRBPs with potential roles in RNA compartmentalization or in anchoring cellular reactions in particular locations. Although membrane-associated proteins have been previously identified as RBPs^30^, there have not been systematic efforts at characterizing memRBPs due to the difficulty of purification using conventional methodologies. Moreover, we observed cRBPs involved in RNA metabolism-related pathways across unexpected cellular locations. This included multiple RNA processing factors that display dual nuclear and cytoplasmic localization in our coRIC map but can only be detected in the nucleus in HPA^20^. On one hand, this reinforces the relevance of our approach for studying intercompartmental crosstalk that may be missed with conventional imaging approaches. On the other, it reminds us that man-made analytical tools such as GO only portray our current knowledge (and bias) of biological processes and will need to be refined or replaced by data-driven cellular map assemblies like MuSIC^48^ or coRIC. Furthermore, it highlights that coRIC captures RBPs in specific locations irrespective of their relative abundance in different compartments. This has implications for the study of hitherto neglected low-expressed proteins acting as RBPs^83,84^. In addition, we observed the enrichment of multiple cRBPs in atypical cellular pathways. Notably, these RBPs have similar features compared to cRBPs in classical pathways, supporting that they may exert similar functions. Consistently, our PPI pathway analysis suggests that RBPs in atypical pathways are hubs for proteins in the same or other pathways, acting as either linkers to RNA metabolism-related processors or scaffolds that stabilize protein complexes. We have also integrated our RBP map with PPI networks from BioPlex^51^ using a machine learning approach to dissect RBP-containing complexes at multiple scales across the cell^50^. This is valuable to identify potential new roles for multifaceted RBPs. For example, we observed that IGF2BP1 is a potential RNA splicing regulator, whereas EIF2A and LARP4 may be implicated in mitochondrial protein translation. Our hierarchical network can offer valuable insights into the role of RNA in mediating RBP complex formation and disassociation by comparing PPI datasets in the presence or absence of RNase.

A major aspect of RBP function related to human disease is how their subcellular localization and interaction with RNA become disturbed under stress. By exploring the effects of *C9ORF72* polyGA aggregates associated to ALS/FTD^66^ and DNA damage, we provide a proof-of-principle that coRIC can be used to gain new insights into disease mechanisms. We found significant RBP subcellular redistribution and RBP complex disruption caused by transient polyGA induction, suggesting a connection between disrupted RNA metabolism and cellular toxicity of *C9ORF72* DPRs *in vivo*. It will be interesting to study how these alterations coordinate with other pathological mechanisms of the *C9ORF72* repeat expansion such as the formation of nuclear RNA foci^85^. The coRIC map also sheds light on DNA damage stress responses. We have identified nearly a hundred *de novo* RBPs in response to DNA damage caused by etoposide treatment. The changed RBPs were enriched in chromatin regulators including transcription factors, expanding the number of DRBPs involved in DNA damage regulation^75^. The latter is consistent with the role of RNA-protein interactions in chromatin dynamics^86,87^, an area of profound significance in multiple other contexts including cancer^88,89^. Our findings emphasize the value of studying RBPs upon stress in specific locations rather than at the whole cell level.

To enable the interactive study of our datasets and pipelines, we have developed a fully searchable online resource (https://db.cngb.org/stomics/rbp/). This will facilitate uncovering new functions and associations, which should be supported by other experimental approaches. As we continue to refine our integrated multimodal framework across topological cellular domains and pathways, several avenues warrant future exploration. First, broadening the datasets to include additional compartments (*e.g.*, multiple chromatin domains) and PPI networks with or without

RNase treatment will enhance our understanding of RBP roles across the multiple scales within the cell. Second, sequencing of compartment-specific RBP-protected footprints will help identify the protein-occupied cis-acting RNA elements, which is critical for studying RBP-mediated compartmentalized RNA regulation. This can be achieved by using partial RNase digestion with streptavidin enrichment of the AGPC-purified interface containing RNP complexes. Third, combination of our methodology and other dynamic tracking methods such as TransitID^90^, will further facilitate the understanding of RBP behavior in response to environmental stimuli or genetic perturbation. In this regard, mis-localization of many RBPs including TDP-43, FUS, and HNRNPA2B1, is observed in various neurodegenerative diseases^4^. Systematic application of coRIC to cells (e.g., induced pluripotent stem cell-derived neurons)^91,92^ bearing mutations in those neurodegenerative disease-related proteins would have enormous value to design rational therapeutic approaches. Collectively, our efforts will greatly expand our understanding of RNA-protein interactions within the normal and perturbed subcellular space.

## Methods

### Cell culture

HEK293T and HeLa cells were cultured in Dulbecco’s modified Eagle’s medium (DMEM)/high glucose (Corning, cat#10-017-CV) supplemented with 10% fetal bovine serum (NATOCOR, cat#SFBE), 1× GlutaMAX (Gibco, cat#35050061), 1× nonessential amino acids (BasalMedia, cat#S220JV) and 1× penicillin/streptomycin (HyClone, cat#SV30010). Cells were grown in 5% CO2 at 37 ℃ and periodically passaged until they reached ∼90% confluency.

### Plasmid and cell line generation

DNA sequences used for plasmid construction were either amplified from the cDNA of HEK293T cells or purchased from GenScript and WZ Biosciences. APEX2 fusion constructs were obtained either from Addgene or through fusing APEX2 sequence to the N or C terminus of bait protein sequencess and subcloning into pLX304 lentiviral vector. The GA50 DPRs sequence and cRBP candidates validated by PAR-CLIP were subcloned into a doxycycline-inducible piggyBac vector with a GFP and a FLAG tag, respectively. The laser microirradiation -validated candidates were subcloned into pKD lentiviral vector with a GFP tag. shRNA oligos targeting *IGF2BP1* were subcloned into pLKO.1 lentiviral vector. Detailed information is provided in **Supplementary Table 8**. For preparation of lentiviruses, HEK293T cells at 70-80% confluence were transduced with lentiviral vectors containing the proteins of interest or shRNA oligos, the packaging vectors psPAX2 and pMD2G using polyethylenimine (PEI, Polysciences, cat#23966) for 8 hours. Then, 48 hours later the culture medium containing lentiviruses was harvested and passed through a 0.45 μm filter (Millipore, cat#SLHPR33RB). Stable cell lines were generated using either the lentivirus or piggyBac transposon system. Briefly, HEK239T cells at 60-70% confluence were transduced with lentiviruses or the piggyBac vector containing the proteins of interest and the transposase vector pBase for 8 hours. Transduced cells were selected with 10 μg/mL puromycin (InvivoGen, cat#ant-pr-1) or 10 μg/mL blasticidin (InvivoGen, cat#ant-bl-1) for a week.

### APEX labelling

HEK293T cell lines expressing APEX2 fusion constructs were cultured to ∼90% confluency. APEX labelling was initiated by replacing the medium with fresh medium containing 500 μM biotin-phenol (APExBIO, cat#A8011-1000) and incubated in 5% CO2 at 37 ℃ for 30 minutes. Next, H2O2 (Sigma-Aldrich, cat#386790-M) was added to a final concentration of 0.1 mM, followed by gentle agitation for 30 seconds. The reaction was quenched by washing three times with the quenching buffer (containing 5 mM Trolox [Sigma-Aldrich, cat#238813-25G], 10 mM sodium ascorbate [Sigma-Aldrich, cat#A4034-100G], and 10 mM sodium azide in Dulbecco’s phosphate-buffered saline [DPBS, BasalMedia, cat#B210KJ]). Unlabeled controls were processed identically except that the H2O2 was omitted.

### Immunofluorescence microscopy

Cells were seeded and cultured on gelatin-coated coverslips (Assistant, cat#41001112) in a 24-well plate until they reached 80-90% confluence. Cells were subsequently washed with DPBS and fixed using 4% paraformaldehyde (PFA, GenXion, cat#JX0100) at room temperature (RT) for 15 minutes. After washing with DPBS, cells were permeabilized with 0.2% Triton X-100 (Sigma-Aldrich, cat#T8787-250ML) in DPBS, followed by incubation with blocking buffer (3% BSA, 0.1% Tween 20 in DPBS) for 1 hour at RT. After incubation with primary antibodies in the blocking buffer overnight at 4 °C, cells were treated with secondary antibodies for 1.5 hours at RT, followed by DAPI (Sigma, cat#D9542) staining for 10 minutes. Images were taken with a Zeiss

LSM710/800/900 confocal microscope. Detailed information of all antibodies used for immunofluorescence is provided in **Supplementary Table 9.**

### Western blotting and silver staining

Western blotting and silver staining were performed as previously described^93^. In brief, protein samples were subjected to SDS-PAGE gel for separation. For Western blotting, proteins from the gel were transferred onto a PVDF membrane (Millipore, cat#IPVH00010). The membrane was subsequently blocked and incubated with primary and secondary antibodies. The signal was generated with ECL (Advansta, cat#K-12045-D50) and visualized with a FUSION SOLO 4M System (Vilber Lourmat). Silver staining of the separated proteins was performed using a Fast Silver Stain Kit (Beyotime, cat#P0017S). Detailed information of all antibodies used for Western blotting is provided in **Supplementary Table 9**.

### Compartmentalized RNA interactome capture (coRIC)

Two 10 cm-dishes with cells reaching ∼90% confluence were used for each condition including the unlabeled control (negative control) and APEX reaction-labeled cells. After APEX labeling, cells were washed twice with DBPS, the remaining DBPS was removed entirely before UV cross-link. Cells were irradiated with 0.15 J/cm^2^ UV light at 254 nm, scrapped off in 2 mL DBPS per plate, and collected by centrifugation at 500 g for 5 minutes at 4 ℃. Whole cell-RBPs were purified following the previously developed AGPC phase partition method OOPS^10^. In brief, the cell pellet from each plate was lysed in 3 mL TRI reagent (MRC, cat#TR118). Lysates were homogenized by vortexing vigorously and then 600 μL chloroform was added followed by vortexing for 20 seconds, and subsequently centrifuged at 12,000 g for 15 minutes at 4 ℃. The upper aqueous phase (containing free RNAs) and the lower organic phase (containing free proteins) were removed. The interface (containing RNA-RBP adducts) was transferred to a new tube and lysed with 1 mL TRI reagent, followed by two additional rounds of AGPC phase separation. The resulting interface was precipitated with 1 mL of cold methanol by centrifugation at 14,000 g for 10 minutes at 4 ℃. The pellet was washed twice with cold methanol and dissolved in 100 μL TEAB buffer (100 mM Triethylammonium bicarbonate [Sigma-Aldrich, cat#T708], 1% SDS) at 95 ℃ for 5 minutes. The samples were then cooled and digested with 5 μL RNase A/T1 mix (2 mg/mL of RNase A and 5,000 U/mL of RNase T1, Thermo Fisher, cat#EN0551) overnight at 37 ℃. The samples were then homogenized with 1 mL TRI reagent. The final round of AGPC phase separation was carried out in which the RNA-bound proteins were released in the organic phase. The organic phase (∼450 μL) from two plates was collected carefully and combined into a new 15 mL tube, followed by adding 3.6 mL cold acetone and 9 μL of 5 M NaCl^94^. After 20 seconds of vortexing and a 30 minute-incubation at −20 ℃, proteins were precipitated by centrifugation at 14,000 g for 10 minutes at 4 ℃. Subsequently, the pellet was washed twice with cold acetone and dissolved in 100 µL TEAB buffer at 95 ℃ for 5 minutes. After taking 5% supernatant as coRIC input, the remaining volume was diluted 10 times with incubation buffer (20 mM Tris-HCl pH7.5, 150 mM NaCl, 2 M urea, 0.5% SDS, 1 mM EDTA pH8.0, 1 mM DTT, and 1× protease inhibitor cocktail [Roche, cat#4693132001]). Biotinylated RBPs were captured by affinity purification. Briefly, the samples were incubated with 100 µL pre-washed MyOne Streptavidin C1 beads (Thermo Fisher, cat#65602) on a rotator gently for 3 hours at RT. Beads were then washed as follows: once with incubation buffer, twice with wash buffer 1 (20 mM Tris-HCl pH 7.5, 150 mM NaCl and 1% SDS), twice with wash buffer 2 (20 mM Tris-HCl pH 7.5, 150 mM NaCl, 6 M urea and 0.1% SDS), twice with wash buffer 3 (20 mM Tris-HCl pH 7.5, 2 M NaCl and 0.5% IGEPAL CA-630 [Sigma-Aldrich, cat#I8896]), and twice with wash buffer 4 (50 mM Tris-HCl pH 7.5). Each wash was performed on a rotator for 5 minutes at RT. The final samples were split, allocating one-third for silver staining and Western blotting as quality controls and the remaining two-thirds for MS sample preparation.

### MS sample preparation

Proteins from pulldown samples in coRIC and immunoprecipitation were processed into peptides using on-bead digestion as previously described^95^. Briefly, the resulting beads from either coRIC or immunoprecipitation were resuspended with 100 μL elution buffer (2 M urea, 100 mM Tris-HCl pH 7.5, and 10 mM DTT) and incubated on a thermomixer for 20 minutes followed by adding 50 nM IAA (Sigma-Aldrich, cat#I5148) and 10-minute incubation at RT. Samples were then digested with 250 ng trypsin (Promega, cat#V5113) on a thermomixer with 1200 r.p.m. at RT for 2 hours in the dark. The supernatant was transferred to a new tube and the beads were incubated again with 100 μL elution buffer on a thermomixer with 1200 r.p.m at RT for 5 minutes. Supernatants were combined and additional 100 ng trypsin was added for proper digestion at RT overnight in the dark. The next day, the digestion was quenched by adding 10 μL of 10% TFA (Sigma-Aldrich, cat#302031). Total proteome samples were processed using the FASP protocol as described before^96^. Briefly, 10 μg of soluble proteins in TEAB buffer were added onto 30 kDa cutoff filter (Sigma-Aldrich, cat#MRCF0R030) and centrifuged at 11,000 g for 15 minutes at RT. 100 μL 50 mM IAA in 8 M urea was used to alkylate proteins for 15 minutes at RT. The filters were then washed thrice with 8M urea and thrice with 50 mM ammonium bicarbonate (ABC, Sigma-Aldrich, cat#A6141). Next, 100 ng of trypsin in 50 μL of 50 mM ABC was added to the filters. The digestion was performed on a thermomixer overnight at 37 °C. The next day, the peptide solution was collected to a new tube by centrifuged at 14,000 g for 10 minutes at RT. The digestion was quenched by adding 10 μL of 10% TFA. All peptide samples were stored at −80 ℃ until further use for LC-MS/MS analysis.

### LC-MS/MS

For coRIC map generation experiments, peptide samples were desalted and subjected to analysis using an Orbitrap Fusion Lumos mass spectrometer (Thermo Fisher Scientific). For perturbation experiments (either coRIC or total proteome), peptide samples were labeled using TMT (Thermo Scientific, cat# A44520). Multiplexed TMT-labeled samples were combined and desalted using homemade C18 desalting tips. Following this, samples were separated into 3 (coRIC) or 10 (total proteome) fractions using Easy-nLC 1200 (Thermo Fisher Scientific) and analyzed using an Orbitrap Fusion Lumos mass spectrometer.

### MS spectral data processing

Raw data from coRIC map samples and stress-perturbed samples were processed using MaxQuant (v.1.6.0.16) and Proteome Discovery (v.2.1), respectively. Tandem mass spectra data were searched against reviewed human protein sequences from UniProt (https://www.uniprot.org/). In MaxQuant, we set Trypsin/P as the cleavage enzyme, allowing a maximum of two missing cleavages per peptide, carbamidomethylation on cysteine as a fixed modification, and oxidation on methionine and protein N-terminus acetylation as variable modifications. The minimum peptide length was set at seven, and the FDR threshold for protein, peptides, and modifications was set as less than 1%. In Proteome Discovery, we kept MS accuracy at 10 p.p.m., allowing up to two missed cleavage sites, carbamidomethylation of cysteine (including TMT tagging of lysine and peptide N terminus for TMT labeled samples) as a fixed modification, and oxidation of methionine and deamidation of asparagine and glutamine as variable modifications. We only accepted rank 1 peptide identifications of high confidence (FDR < 1%), with a minimum of two high-confidence peptides per protein required for identification. Before proceeding with the statistics, the common contaminant proteins and glycoproteins were removed from the protein-level output files as suggested in OOPS^10^.

### SAINTexpress analysis

For each prey protein enriched by a specific bait in a coRIC experiment, SAINTexpress (v3.6.3) was applied to calculate the probability of being a true interaction by using its ion-mass intensity against the control samples. To obtain RBPs enriched in the adjacent compartments, the protein-level output files from nearby-compartment baits were used as spatial reference and the proteins with Bayesian FDR (BFDR) ≤ 0.05 were defined as enriched RBPs. Specific information of one-on-one comparisons is shown as follows, the mitochondrial-enriched RBPs: MITO-APEX vs NES-APEX; nucleus-enriched RBPs: NLS-APEX vs NES-APEX; nucleolus-enriched RBPs: NIK-APEX vs NLS-APEX; OMM-enriched RBPs: OMM-APEX vs NES-APEX. To obtain whole cell cRBPs, the protein-level output files from all compartment baits and control samples (H2O2 omitted) were used and the proteins with BFDR ≤ 0.01 were defined as cRBPs.

### Assessment of compartment-enriched RBPs

For the assessment of mitochondria-enriched RBPs, human mitochondrial proteins were collected from MitoCarta 3.0 (https://personal.broadinstitute.org/scalvo/MitoCarta3.0/)^97^, and previously annotated RBPs were collected from RBPbase (https://apps.embl.de/rbpbase/)^98^. Known mitochondrial RBPs were defined by intersecting the human mitochondrial proteins with the previously annotated RBPs, whereas known mitochondrial non-RBPs were defined as those human mitochondrial proteins not overlapping with these annotated RBPs. To compare the nucleus-enriched candidate RBPs, curated nucleus RBP datasets were defined by the intersections of the data from two nucleus RBPs studies serIC and eRIC^5,6^. The GO comparison of nucleus-, nucleolus-, and OMM-enriched RBPs from coRIC with APEX-PS was performed using *compareCluster* function with the R package clusterProfiler (v.4.0.5)^99^.

### NMF analysis and *t*-SNE map generation

NMF analysis was performed using the Python module scikit-learn (v.0.18.1) as previously described^21^. In brief, the dimensions of our input coRIC matrix were 1,768 × 22, in which 1,768 preys from 22 baits passed a BFDR ≤ 0.01 cut-off. Prey intensity had their average value in the control subtracted and was rescaled from 0 to 1 across baits to represent the signals with equal weight. Next, NMF was run on this matrix with different numbers of ranks ranging from 2 to 22 (maximal size of baits). For each NMF run, the most abundant prey in each rank was annotated by GO cellular compartment using R package gProfiler2 (v.0.2.2)^100^. The optimal number of ranks to use for NMF was determined by maximizing the number of preys assigned to known localizations and minimizing the overlapping GO terms between ranks. The NMF run with 14 ranks was outperformed and a representative GO term or terms were chosen to define the rank for further analysis. The coRIC map was then built from the NMF ***W*** matrix corresponding to 14 ranks using *t*-SNE.

### Comparison of cRBPs with RBPome and protein localization datasets

To define novel RBPs among the cRBPs, we first generated a curated RBP list using phase separation-based RBP datasets from HEK293 cells (OOPS, XRNAX, and PTex)^10,18,19^ together with proteins annotated as ‘RNA binding’ in the GO database. Those RBPs that did not appear in any of the four databases were classified as novel RBPs. To evaluate the localization specificity of cRBPs, we downloaded the subcellular localization data from HPA (v.21; https://www.proteinatlas.org/)^20^ and HCM^21^ (v.1; https://cell-map.org/). Fisher’s exact test was performed to assess the enrichment of cRBPs in those compartments defined by HPA and HCM. The −log10 adjusted *P* value served to indicate the relative concurrence between the localization of cRBPs and the compartments identified by HPA and HCM.

### Analysis of cRBP characteristics

The human proteome-wide biophysical properties, predicted RNA-binding probability score, and experimentally identified PTM sites were obtained from published datasets^28,53,101^. The PTM density of each protein was calculated by dividing the number of experimentally-identified sites by the amino acid length. And the protein domains of cRBPs were annotated using Pfam (https://www.ebi.ac.uk/interpro/entry/pfam/) or UniProt (https://www.uniprot.org/)^102,103^. All proteins contained in the GO term/terms defined by NMF rank were used to compare the characteristics and PTM density of cRBPs and proteome in the corresponding compartments. The value of each feature and PTM density was average and scaled for visualization. Fisher’s exact test and standard FDR control method were used to determine the protein domains enriched in each compartment relative to the total proteome. Transmembrane proteins were obtained using the UniProt API with ‘types=TRANSMEM’, and the memRBPs were defined by intersecting different RBP profiling datasets with the transmembrane proteins.

### Subcellular RBP pathway analysis

To elucidate the biological pathways in which cRBPs participate, cRBPs (including multi-localizations) in each compartment analyzed using the *enrichGO* function with the R package clusterProfiler (v. 4.0.5). The pathways with −log10 adjusted *P* value ≥ 2 and log2 fold-enrichment (compartment vs genome-wide) ≥ 1 were considered as significantly enriched terms.

### Pathway-RBP interaction score

The score was generated to scale the relative RBP interaction capacity among the enriched proteins in each atypical pathway. The PPI dataset was obtained from BioPlex 3.0 (https://bioplex.hms.harvard.edu/)^51^. previously annotated RBPs collected from RBPbase^98^ were defined as canonical RBPs and the remaining proteins that were identified by more than 2 human RIC studies were defined as non-canonical RBPs. To avoid self-validation and occasional cases, the cRBP-interacting proteins that belong to canonical RBPs were excluded, the remaining proteins with ≥ 4 interactions and the pathways containing ≥ 4 proteins after filtering were used for later analysis. The RBP interaction score (RIS) of protein was first defined by determining the proportion of canonical RBPs that each protein interacts with. The pathway-RBP interaction score (PRIS) is derived from the sum of the RIS for each protein in the pathway divided by the total number of proteins.

### RBP hierarchical network generation

We followed the model of a previous report^48^. In brief, the overlapping 1,465 proteins from coRIC and BioPlex 3.0 dataset^51^ were used for integration. To generate an embedding vector of the coRIC dataset, we first performed Pearson correlation analysis of the 1,465 proteins using the NMF ***W*** matrix in the coRIC map. The resulting protein pair with Pearson correlation ≥ 0.9 were retained. Next, the protein pair matrix from either coRIC or the BioPlex dataset was embedded with the node2vec method from PecanPy (v.2.0.8) into a 1,024-dimensional vector separately^104^. The 1,465-protein list was used to calculate the gold standard protein-protein proximity based on the Cellular Component branch of the GO database. The generated curated protein pair distances were used as training labels, and the embedded matrix was integrated and constructed with random forest regressors to predict the pairwise distance of any protein directly from its data embeddings. The generated protein-pair distance matrix for all proteins was used for pan-resolution community detection using CliXO (v.1.0) with different parameters. α reflects the step size by which network stringency (protein pair distance) is lowered for progressive cycles of clique detection; a smaller α tends to generate a deeper hierarchy, in which the differences between parent and child communities tend to be smaller. β reflects the stringency of merging overlapping cliques; a higher β tends to merge cliques less frequently, resulting in a broader hierarchy with more sibling systems with larger overlaps; *m* reflects the modularity of each community, a higher *m* reduces the number of communities. To determine which combination of parameters had better performance, we calculated the known protein pairs from two independent protein-interaction datasets, CORUM^52^ and HCM^21^, that were recovered by different parameter-generated hierarchies. The goal was to generate the simplest hierarchical network with most of the known protein pairs recovered. Finally, the parameters (α = 0.01, β = 0.5, and *m* = 0.002) were selected.

### Network visualization

All PPI and hierarchical networks were visualized using Cytoscape (v.3.9.1)^105^, except for the PPI networks of memRBPs, which were generated using the *MCODE* function of Metascape (https://metascape.org/gp/index.html)^106^.

### PAR-CLIP

PAR-CLIP validation of cRBP was performed as previously described^107^. Briefly, HEK239T cells were transduced with plasmids expressing the FLAG-tagged proteins of interest. Then, 24 hours later, they were treated with 0.2 mM 4sU (Sigma-Aldrich, cat#T4509-25MG) and 1μg/mL doxycycline (Sigma-Aldrich, cat#D9891-1G) for 16 and 24 hours, respectively. Cells were cross-linked with 0.4 J/cm^2^ of 365 nm UV light and lysed in NP40 lysis buffer (50 mM Tris-HCl pH 7.5, 100 mM NaCl, 2 mM EDTA, 1% NP40, 0.1%SDS, 0.5 mM DTT, 40 U RNase inhibitor [New England Biolabs, cat#M0314L], and 1× protease inhibitor cocktail [Roche, cat#4693132001]). Lysates were treated with 1 U/mL RNase T1 (Sigma-Aldrich, cat#R1003) for 15 minutes at 22 ℃. Pre-washed magnetic beads conjugated with FLAG antibodies (Sigma-Aldrich, cat#M8823) were added to the lysate and incubated for 3 hours at 4 ℃. Beads were washed three times with CLIP washing buffer (50 mM Tris-HCl pH 7.5, 300 mM NaCl, 0.05% NP40 and 0.5 mM DTT), followed by treatment with 10 U/μL RNase T1 in CLIP washing buffer for 15 minutes at 22 °C. Beads were next washed three times with dephosphorylation buffer (50 mM Tris-HCl pH 7.5, 100 mM NaCl, 10 mM MgCl2 and 1 mM DTT) and treated with 0.5 U/mL calf intestinal alkaline phosphatase (New England Biolabs, cat#M0290) in dephosphorylation buffer for 10 minutes at 37 ℃. The RBP-bound RNAs were subjected to on-bead biotinylation using Pierce RNA 3’ End Biotinylation Kit (Thermo Fisher, cat#20160). After labeling, the beads were washed three times with CLIP washing buffer, resuspended with 1×NUPAGE LDS buffer (Invitrogen, cat#NP0007) and heated at 95 °C for 5 minutes to elute the RNP complexes. The elution was separated with Novex Bis-Tris 4-12% PAGE gel (Invitrogen, cat#NP0323BOX) and transferred onto a nitrocellulose membrane (Cytiva, cat#10600001) for Western blotting or for RNA detection using a Chemiluminescent Nucleic Acid Detection Module Kit (Thermo Fisher Scientific, cat#89880).

### Immunoprecipitation (IP) and AP-MS analysis

Twenty million HEK293T cells were lysed in 1 mL TNE lysis buffer (50 mM Tris HCl pH 7.5, 250 mM NaCl, 0.5% IGEPAL CA-630, 1 mM EDTA, and 1× protease inhibitor cocktail). The lysate was incubated on a rotator for 30 minutes at 4 ℃, followed by centrifugation at 14,000 g for 10 minutes at 4 ℃. The supernatant was transferred to a new tube and diluted with 1 mL TNE dilution buffer (50 mM Tris HCl pH 7.5, 50 mM NaCl, 0.5% IGEPAL CA-630, 1mM EDTA, RNase inhibitor and 1×protease inhibitor cocktail). 2 µg endogenous antibodies were added to the diluted lysate and incubated on a rotator overnight at 4 ℃. Next day, 20 µL of pre-washed Protein A/G magnetic beads mixture (Invitrogen, cat#10002D and cat#10004D) was added to the samples and incubated on a rotator for another 2 hours at 4 ℃. Beads were washed twice with TNE wash buffer (50 mM Tris HCl pH 7.5, 150 mM NaCl, 0.5% IGEPAL CA-630 and 1 mM EDTA), then resuspended in the same buffer and evenly split into two tubes. Next, half of the beads were treated with 2 µg BSA (BSA-treated group), and the other half were treated with 2 µg RNase A (Thermo Fisher, cat#EN0531) (RNase-treated group), on a thermomixer at 1000 r.p.m for 30 minutes. The treated beads were then washed twice with TNE wash buffer and processed for MS analysis as in the MS sample preparation described above. Differentially-bound RBP analysis was performed using the R package DEP (v.3.17) as previously described^108^. Briefly, the protein-level output files from each antibody were used, and the interactors of each protein were obtained by comparing each antibody with the IgG sample. RNA-dependent interactors are obtained by comparing the BSA-treated with RNase-treated sample from the same antibody. The interactors of each protein were compared with the reported interactors from the integrated PPI dataset in BioGRID (https://thebiogrid.org/)^109^. The remaining proteins that do not overlap with known interactors were defined as newly identified interactors.

### RNA-seq data processing, differential splicing analysis and RBP-bound motif analysis

Total RNA from HEK293T cell lines expressing shRNA targeting *luciferase* (sh*Luc*) or *IGF2BP1* (sh*IGF2BP1*) were extracted using TRIzol. RNA libraries were constructed using the ribosomal RNA deletion method and then sequenced on the DIPSEQ platform, generating ∼80 million 100 bp paired-end reads per sample. The raw sequence files were trimmed of adapters and filtered low-quality reads using fastp (v.0.20.1). Reads were aligned to the hg38 reference genome using STAR (v.2.7.9a), followed by PCR duplicate depletion using sambamba (v.0.8.8). The aligned reads were assigned with GRCh38.p13 genome annotation from GENCODE using featureCounts (v.2.0.2). rMATs (v.4.1.2) was used to quantify the changes in five different types of splicing events in sh*IGF2BP1* relative to sh*Luc* controls across three biological replicates. Statistically significant splicing changes were selected based on the criterion inclusion level ≥ 10% and FDR ≤ 0.01. Sashimi plots for visualizing splicing changes were generated using rmats2sashimiplot (v. 2.0.4). Detailed information on these splicing changes is listed in **Supplementary Table 6**. For analysis of the enriched RBP motifs related to the skipping exon (SE) events, the human RBP motifs matrix was downloaded from the oRNAment database (https://rnabiology.ircm.qc.ca/oRNAment)^110^ and converted to HOMER motif format using the *write_homer* function from the R package universalmotif (v.1.12.4). To assess the enrichment of human RBP motifs in the regions of SE events, we first extracted 300 bp flanking region sequences upstream and downstream of the cassette exons as target sequences and randomly selected sequences from the reference genome as background sequences using findMotifsGenome.pl in Homer (v.4.9.1) with the parameter ‘hg38 - size given -rna -len 7 -dumpFasta’. The target sequences were further used for calculating known RBP motif enrichment compared with background sequences using the AME module in the MEME suite (v.5.5.2)^111^. Fisher’s exact test was performed and the *P* values were adjusted using a Bonferroni correction. For visualization, the RBP motif from IGF2BP1 interacting proteins with the adjusted *P*-value ≤ 0.001 was chosen.

### Semi-quantitative PCR

We used cDNA from HEK293T cell lines expressing shRNA targeting *luciferase* (sh*Luc*) or *IGF2BP1* (sh*IGFBPP1*). Primers were designed to flank the region of the cassette exons and ensure that the length of PCR amplicons was less than 500 bp. PCR products were separated by 3% low melting agarose gel (Lonza, cat#50111) and detected with Gel Image System (Tanon 1600). The percentage of exon-inclusion of tested isoforms was calculated using the PCR amplicon signal measured in ImageJ.

### Stochastic Optical Reconstruction Microscopy (STORM)

0.1 million HEK293T or HeLa cells were seeded on borosilicate glass bottom 8-well chambers (µSlide Ibidi, cat#80827) and cultured for 24 hours. Cells were subsequently washed with DPBS and fixed using 4% PFA and 0.1% glutaraldehyde (Sigma-Aldrich, cat#G5882-100ML) in DPBS for 10 minutes at RT. After washing with DPBS, cells were incubated with 1 mg/mL NaBH4 (Innochem, cat#A58223) for 7 minutes at RT and then washed with DPBS. Cells were then blocked with STORM blocking buffer (3% BSA and 0.1% Triton X-100 in DPBS) for 10 minutes at RT, followed by incubation with primary antibodies in STORM blocking buffer overnight at 4°C. Negative control samples were prepared in parallel except the primary antibody was omitted. After washing with 1% BSA in DPBS, cells were incubated with secondary antibodies for 1 hour at RT, followed by DAPI staining for 10 minutes. Images were acquired on Nanoimager S Mark II (Oxford Nanoimaging). Before proceeding to imaging, freshly prepared imaging buffer (100 mM Cysteamine MEA [Sigma-Aldrich, cat#30070], 5% Glucose [Sigma-Aldrich, cat#G8270], 0.5 mg/mL glucose oxidase [Sigma-Aldrich, cat#G2133], and 40 mg/mL catalase [Sigma-Aldrich, cat#C100] in DPBS) was added in the well. 488 nm and 647 nm lasers were used as excitation at constant 100% and 70% laser power, respectively. Simultaneous image acquisition was performed with a 10 ms exposure time for 45,000 frames. Raw images were processed in ONI NimOS (v.1.19.4). Background signals were corrected by using the *Filters* function (Localization precision X and Y: 10∼30 nm; Sigma X and Y: 20∼250 nm; *p*-value: 0.99∼1). The localization lists of each image were exported and converted into bin files using custom-built software in MATLAB 2015a^112^. The generated bin files were further processed using the AFIB-Plugin in ImageJ (v.1.50i). To determine the mitochondrial localization density of each protein, we first defined the mitochondrial regions by outlining the signal derived from the outer mitochondrial membrane marker TOMM20. The mitochondrial localization density was further calculated by dividing the signal of each protein that localized in the mitochondria by the area of the mitochondrial regions. Detailed information of all antibodies used for STORM is provided in **Supplementary Table 9.**

### Total and subcellular RNA-binding proteome preparation under stress

For polyGA DPRs-induced stress, HEK293T cell lines expressing NES-APEX, NLS-APEX, NIK-APEX, or MITO-APEX were transduced with a piggyBac vector containing GFP-GA50 and the transposase vector pBase to generate co-expression cell lines. The resulting cell lines were expanded and cultured with medium supplemented with 1 μg/mL doxycycline for 24 hours, while control cells remained untreated. For DNA double-strand break stress. HEK293T cell lines expressing NLS-APEX were treated with 50 μM etoposide (TargetMol, cat#T0132) for 1, 6 or 12 hours, while control cells remained untreated. Two 10 cm-dishes with ∼90% confluent cells were used for each condition. Cells from each condition were subjected to RBP purification using coRIC as described above. For total proteomes, 100 μL TRI reagent homogenized lysate from each sample was taken and the proteins were precipitated by adding 900 μL methanol and centrifuged at 14,000 g for 10 minutes at 4 ℃. The pellet was washed twice with cold methanol, followed by dissolution in 50 μL TEAB buffer for 10 minutes at 95 ℃. Samples were further processed as in the MS sample preparation described above.

### Quantification of differential RNA-binding and proteome upon stress

Protein abundance was log2 transformed and center-median normalized within each sample. To determine proteins with a change in abundance or RNA binding, we used the *lm* function in the R package limma (v.3.48.3). Briefly, protein abundance was modeled as a function of the treatment or timepoint for the dipeptide and DNA damage experiments, respectively. The moderated *t*-test was performed, and the proteins with Benjamini-Hochberg adjusted *P* value ≤ 0.01 (polyGA vs ctrl for the dipeptide experiment, and each timepoint [1, 6, and 12 hours] versus 0 hour for DNA damage stress) were considered to have significant changes in abundance or RNA-binding. To remove the potential background produced from the APEX reaction in polyGA samples, the significantly changed RBPs that overlapped with the coRIC map were selected for further analysis. In addition, to eliminate the background and retain *de novo* RBPs in DNA damage samples, the altered RBPs that overlapped with the nuclear compartments in the coRIC map and HPA were selected for further analysis. To analyze the pattern of the changed proteins upon DNA damage, we used hierarchical clustering with the *hclust* function in R. Detailed information on the differentially expressed proteins and differentially-bound RBPs are provided in **Supplementary Table 7**.

### Laser microirradiation-induced DNA damage stress

Laser microirradiation was performed as previously described^79^. Briefly, HeLa cells were grown on a dish with a thin glass bottom (NEST, cat#801006) and then locally irradiated with a 365 nm pulsed nitrogen UV laser (16 Hz pulse, 55% laser output) generated from a Micropoint System (Andor). Images were captured in real time every 20 seconds for 10 minutes under a DragonFly confocal imaging system (Andor).

#### Acknowledgments

We thank all other members of our laboratories for their support; L. Liu, C. Liu, and X. Xu (BGI Research) for constructive suggestion on bioinformatic analyses; X. Wei and T. Yang (China National GeneBank) for construction of online database; X. Liu (Guangzhou Institutes of Biomedicine and Health) for kindly sharing anti-TOMM20 antibody; R. Shi (Guangzhou Institutes of Biomedicine and Health) for the assistance of MS measurement; M. Vermeulen (Radboud University) for the suggestions of proteomics design and analysis; and X.D Fu (Westlake University) for constructive suggestions on the manuscript. This work was supported by the National Natural Science Foundation of China (32201214 to Y.Lv, 31900617 to X.G., 22004021 to M.S., 31971177 to M.P.C., 32090031 to X.X., 32211530050 to F.A. and M.A.E., and U20A2015 to M.A.E.); the Guangdong Basic and Applied Basic Research Foundation (2023A1515010839 to Y.Lv, c23140500002783 to S.K., 2021B1515120075 to M.A.E.); the Guangzhou Science and Technology Foundation (2023A04J0728 to Y.Lv); the Guangdong Pearl River Talent Program (2019QN01Y051 to X.Z.); the Science and Technology Planning Project of Guangdong Province (2020B1212060052 to X.Z.). X.G. is supported by the CRG-GDL international PhD programme between the Foundation Centre for Genomic Regulation and the Bioland Laboratory.

## Author contributions

Conceptualization: Y.Lv, X.G., M.A.E. Methodology: Y.Lv, X.G., J.H., S.K., J.Y., M.T., M.J. Investigation: Y.Lv, X.G., J.H., S.K., J.Y., M.T., J.Z., M.S., Y.Lu, J.W., A.W., A.C.G., X.Z., B.P., X.X. Visualization: Y.Lv, X.G., J.H., J.Y., A.C.G. Funding acquisition: Y.Lv, X.G., S.K., M.S., M.P.C., X.X., F.A., X.Z., M.A.E. Project administration: Y.Lv, X.G., D.W., M.P.C., L.D.C., X.Z., M.A.E. Supervision: Y.Lv, M.A.E. Writing – original draft: Y.Lv, M.A.E. Writing – review & editing: Y.Lv, X.G., J.H., S.K., M.T., G.V., M.A.M., W.L., L.W., Y.Lai, P.H.M., E.L., J.M., A.P.H, M.P.C., Y.Z., M.A.E.

---

**Extended Data Fig. 1.**
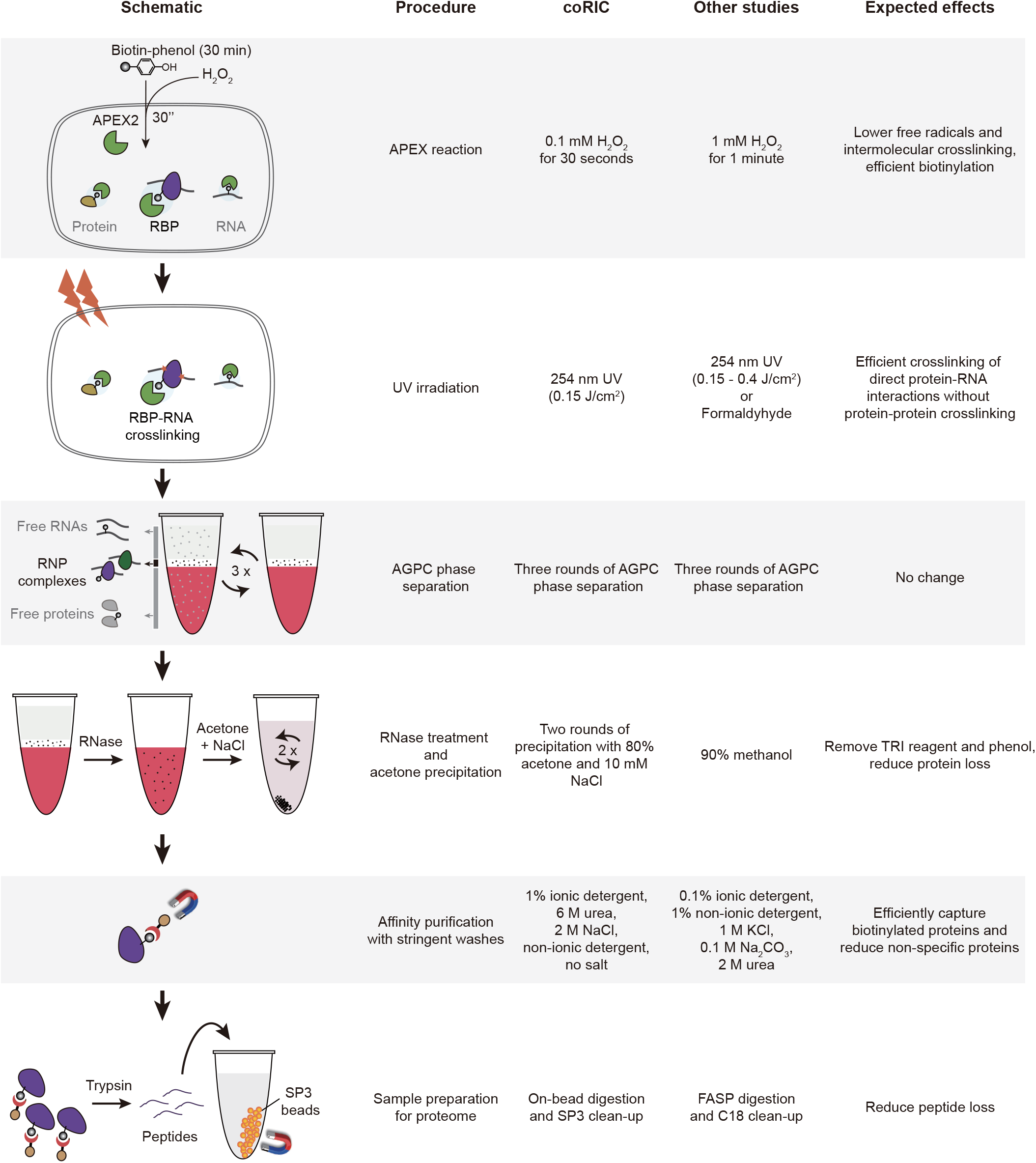
Schematic representation of the optimized procedures in coRIC (related to Fig. 1). The establishment of the coRIC technique required optimizing several key steps. These included: reducing the concentration of reaction catalysts and reaction times for APEX to minimize cross-linking events induced by free radicals^115^; recovering proteins by repeated precipitation with acetone^94^ to fully remove TRI reagent and phenol, which are incompatible with mass spectrometry; introducing high salt-concentration buffers during the washing process to remove non-specific background; and employing on-bead digestion and SP3 desalting strategies to minimize peptide loss^116^ (see Materials and Methods). RNP, ribonucleoprotein; AGPC, acidic guanidinium thiocyanate-phenol-chloroform; SP3, solid-phase-enhanced sample preparation, FASP, filter aided sample preparation.

**Extended Data Fig. 2.**
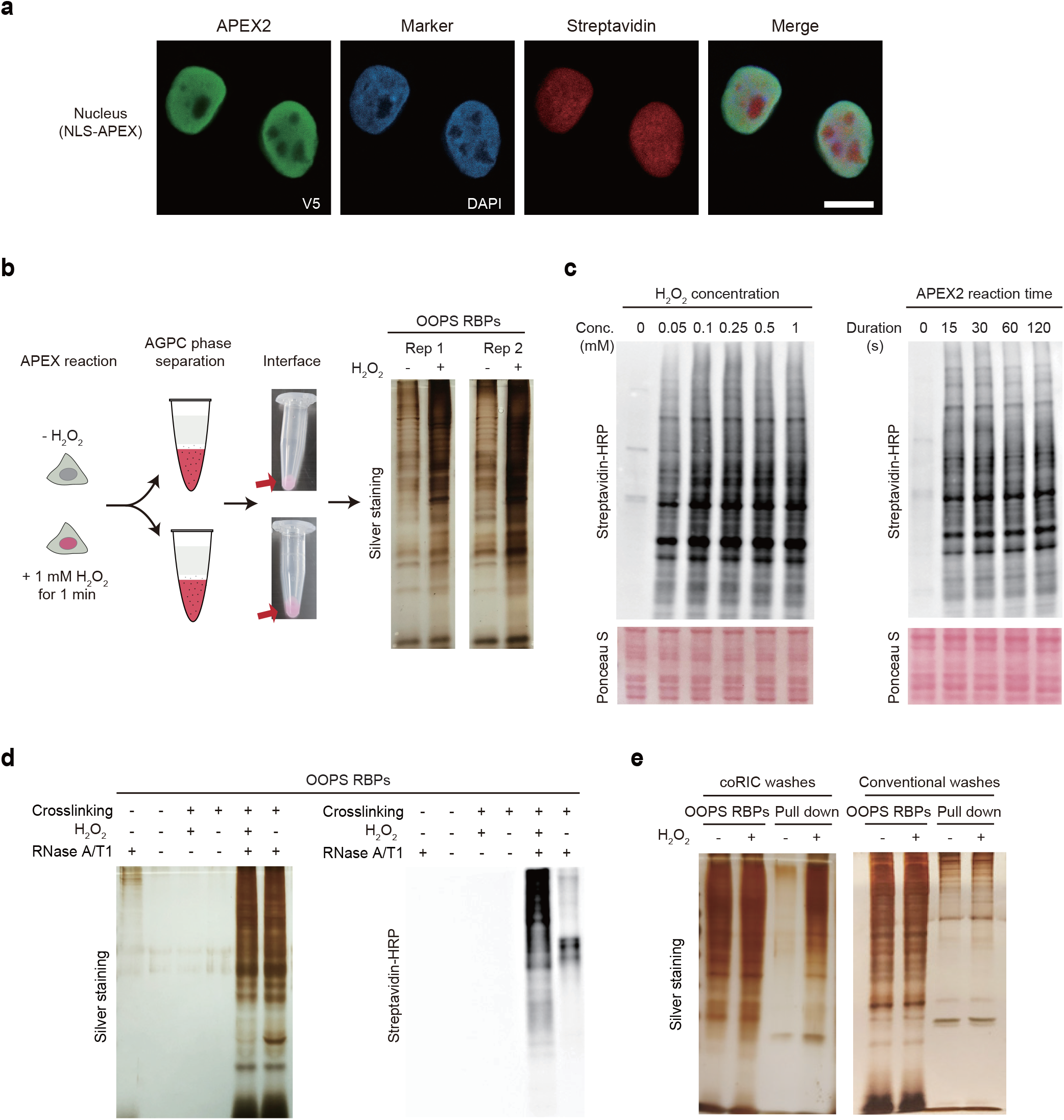
Establishment of coRIC using NLS-APEX cell line (related to Fig. 1). **a,** Confocal immunofluorescence of NLS-APEX and biotinylation signal after APEX reaction. NLS-APEX fusion protein was visualized using V5 tag staining (green). DAPI stained nuclei. Biotinylation activity was detected by staining with streptavidin-Alexa Fluor 647 (red). Scale bar, 10 μm. **b,** Schematic representation of the increasing volume of the interface using a conventional APEX reaction condition (1 mM H2O2 for 1 minute). The protein abundance of the interface, representing orthogonal organic phase separation-purified RBPs (OOPS RBPs), was analyzed using silver staining. **c,** Optimization of APEX reaction for coRIC. Cells were subjected to APEX reaction with different H2O2 concentrations (left panels) and reaction times (right panels), then lysed and analyzed by staining with streptavidin-HRP or Ponceau S. The optimized condition (0.1 mM H2O2, 30 seconds), in which produced comparable biotinylation signal than conventional condition, was chosen for all subsequent experiments. **d,** Validation of the optimized APEX reaction condition using OOPS RBPs. Silver staining (left) and streptavidin-HRP blot (right) analysis demonstrated efficient biotinylation in OOPS RBPs without increasing their protein abundance using the optimized APEX reaction. **e,** Optimization of the washing conditions used for affinity purification. Silver staining analysis of the improvement of pulldown specificity with the washing condition used in coRIC compared with a conventional protocol^11^.

**Extended Data Fig. 3.**
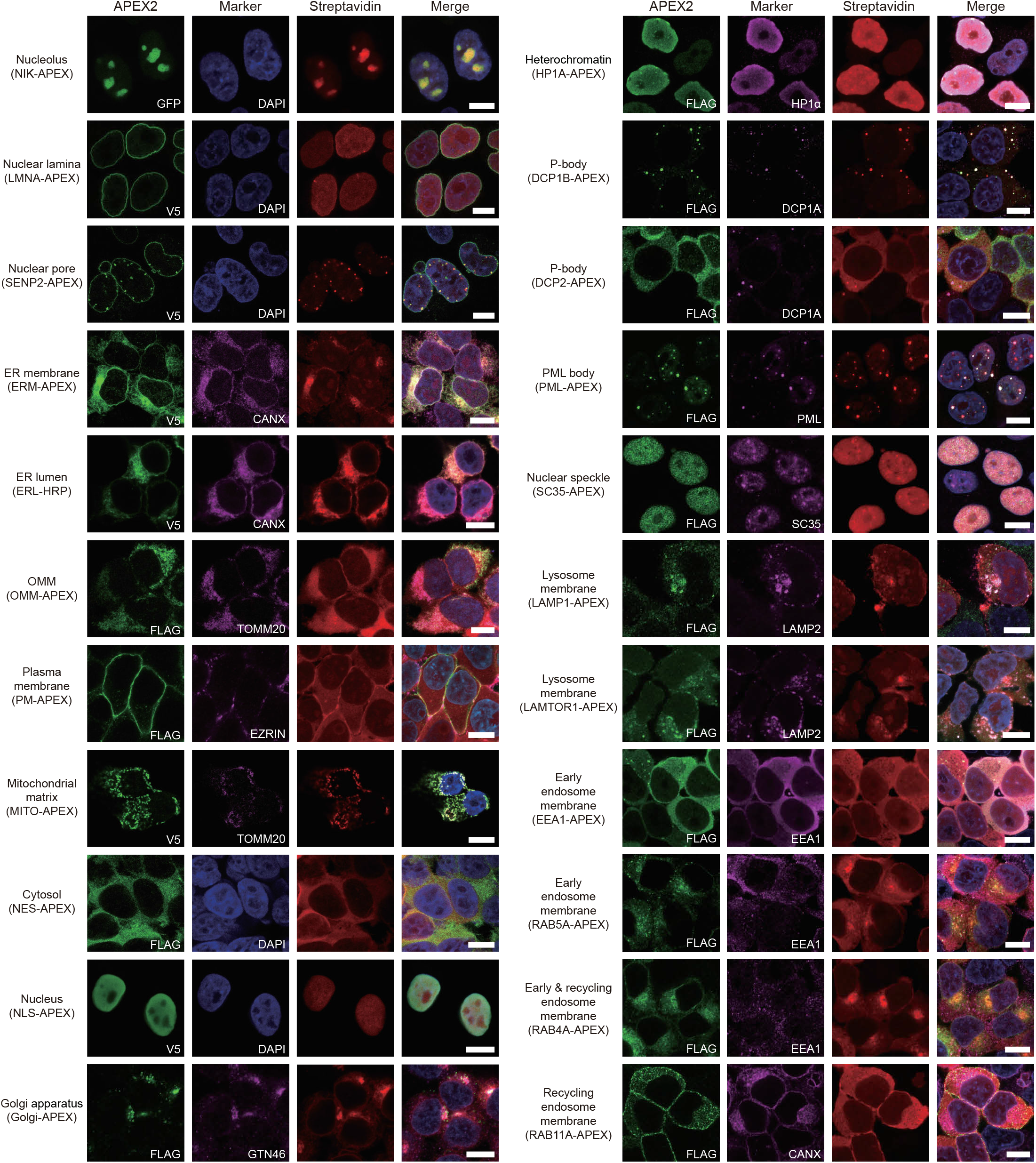
Fluorescence imaging of the 22 bait cell lines used for coRIC (related to Fig. 1). HEK293T cells expressing the APEX2 fusion protein and visualized by immunofluorescence with GFP/V5/FLAG antibody (green). Biotinylation was visualized by streptavidin-Alexa Fluor 647 (red). Antibodies for CANX, TOMM20, EZRIN, TGN46, HP1α, DCP1A, PML, SC35, LAMP2 and EEA1 were used as markers for the ER, mitochondria, plasma membrane, Golgi apparatus, heterochromatin, P-body, PML body, nuclear speckle, lysosome, and early endosome, respectively. DAPI stained nuclei. Scale bars, 10 μm. ER, endoplasmic reticulum; OMM, outer mitochondrial membrane.

**Extended Data Fig. 4.**
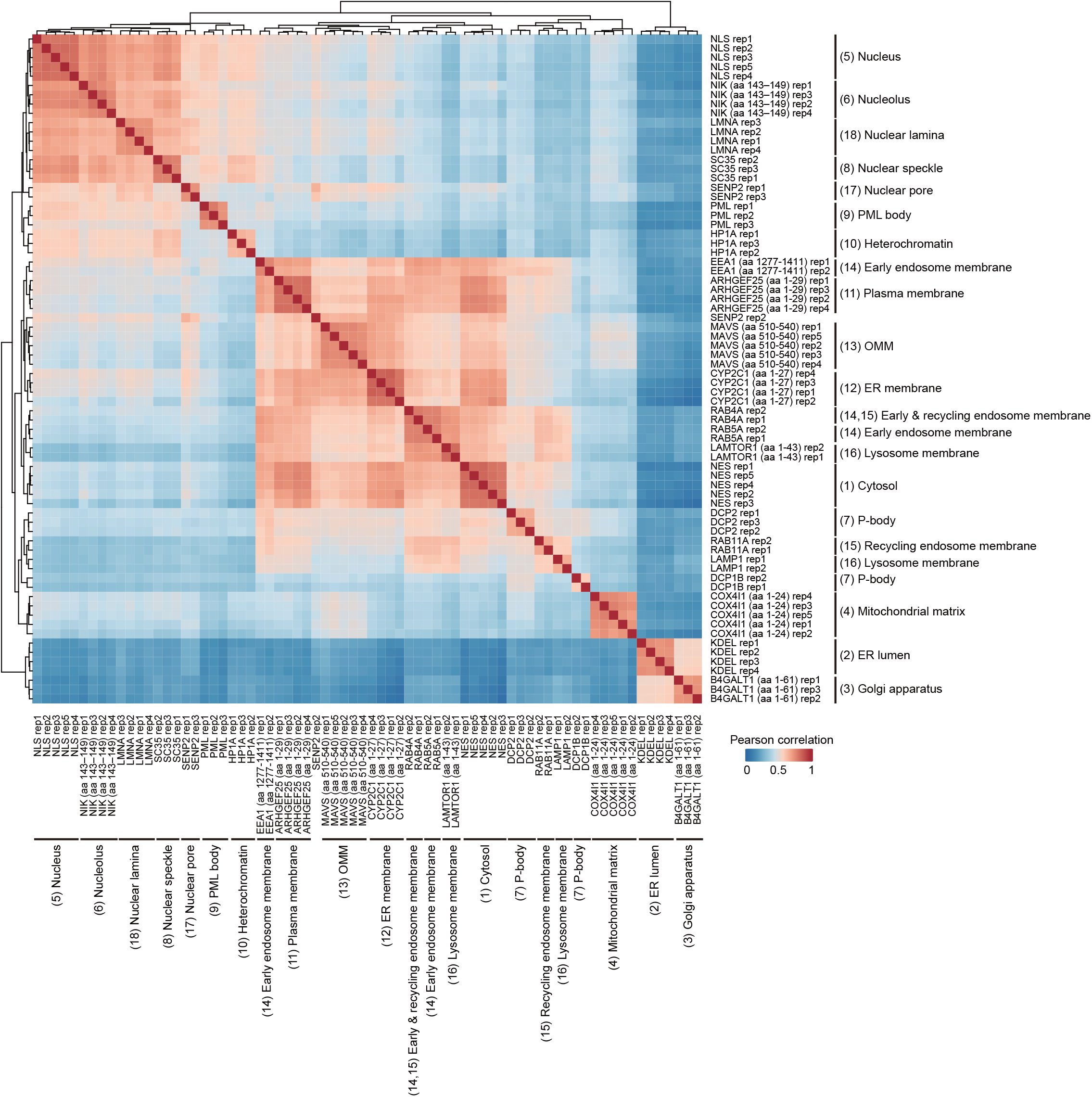
Correlation of coRIC datasets (related to Fig. 1). Heatmap representing the Pearson pair-wise correlation scores of the coRIC datasets including biological repeats for each bait.

**Extended Data Fig. 5.**
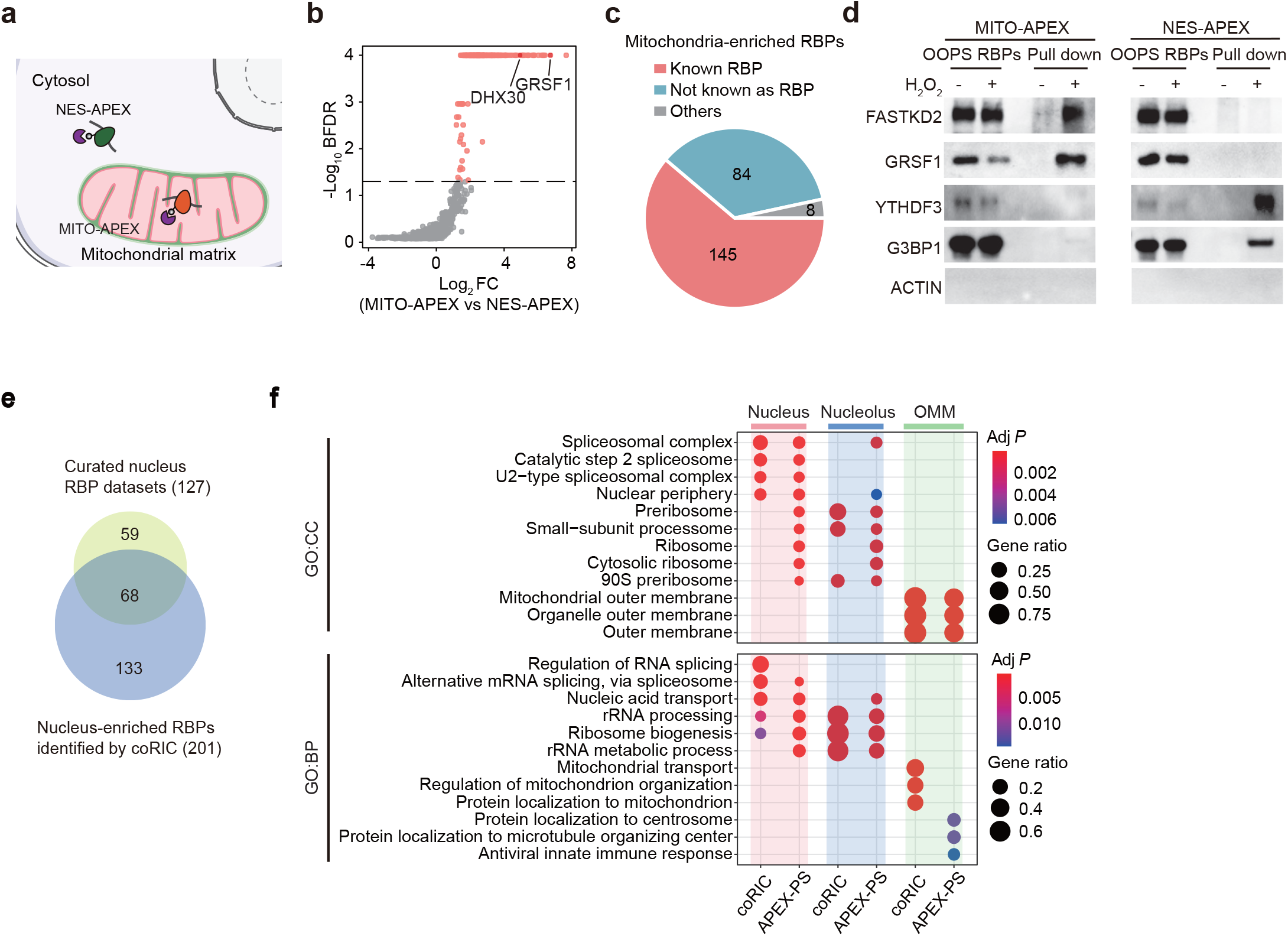
Identification of compartment-enriched RBPs by coRIC (related to Fig. 1). **a,** Schematic showing APEX labelling of RNP complexes in MITO-APEX (mitochondrial matrix) and NES-APEX (cytosol). **b,** SAINTexpress scoring of the coRIC datasets from MITO-APEX cell line versus the neighboring spatial control NES-APEX. Proteins with a Bayesian false discovery rates (BFDR) of zero were given a value of 0.0001 for plotting. Proteins with the BFDR ≤ 0.05 are defined as mitochondria-enriched candidate RBPs and colored in pink. Two canonical mitochondrial RBPs GRSF1 and DHX30 are highlighted in red. Log2 FC, log2 fold-change. **c,** Pie chart showing the proportion of different categories of 236 mitochondria-enriched RBPs. Known mitochondrial RBPs are colored in red, known mitochondrial proteins without prior RBP annotation are colored in light blue, and others are colored in grey (see Materials and Methods). **d,** Western blot validation of selected RBPs captured by coRIC in MITO-APEX and NES-APEX cell lines. ACTIN is the negative control. **e,** Venn diagram comparing the nucleus-enriched RBPs identified by coRIC and curated nucleus RBPs datasets from other studies^5,6^ (see Materials and Methods). **f,** Comparison of the RBPs enriched in the nucleus, nucleolus, and OMM from coRIC and APEX-PS method^15^ using GO analysis (Fisher’s exact test, Benjamini-Hochberg corrected *P* ≤ 0.01).

**Extended Data Fig. 6.**
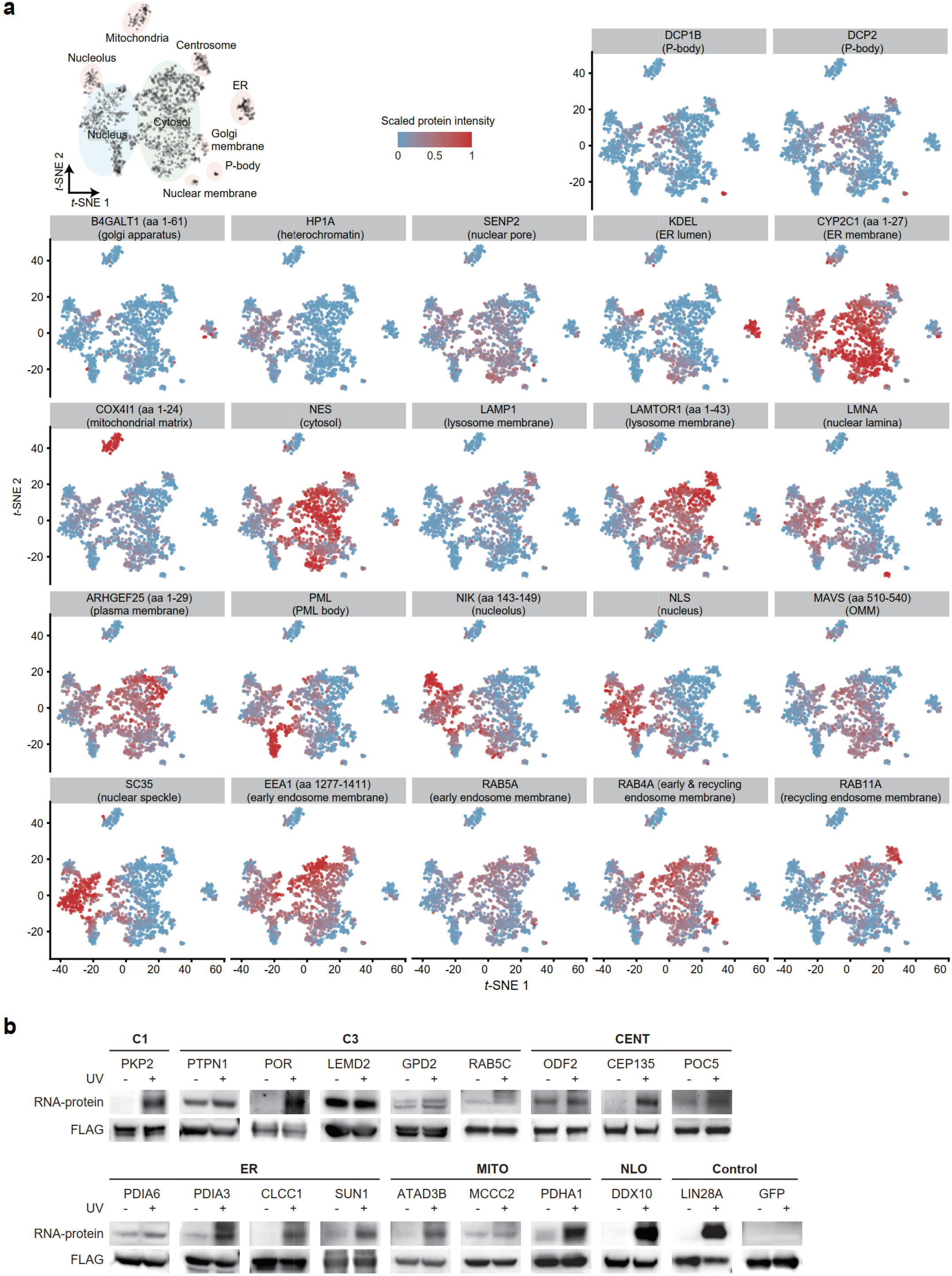
Spatial distribution of cRBP captured by each bait and PAR-CLIP validation (related to Fig. 1). **a,** Projection of cRBPs captured by each bait onto the *t*-SNE map generated in (Fig. 1c). The abundance of cRBPs was scaled from 0 to 1 across different baits to ensure that the signal of the preys are represented with equal weight. The compartments in the original *t*-SNE map are marked with ellipses. *t*-SNE*, t*-distributed stochastic neighboring embedding. **b,** PAR-CLIP validating the RNA-binding activity of 17 selected cRBPs from different compartments isolated by coRIC. FLAG was used to confirm the protein size and equal loading. GFP and LIN28A were used as the negative and positive control, respectively. PAR-CLIP, photoactivatable-ribonucleoside-enhanced crosslinking and immunoprecipitation.

**Extended Data Fig. 7.**
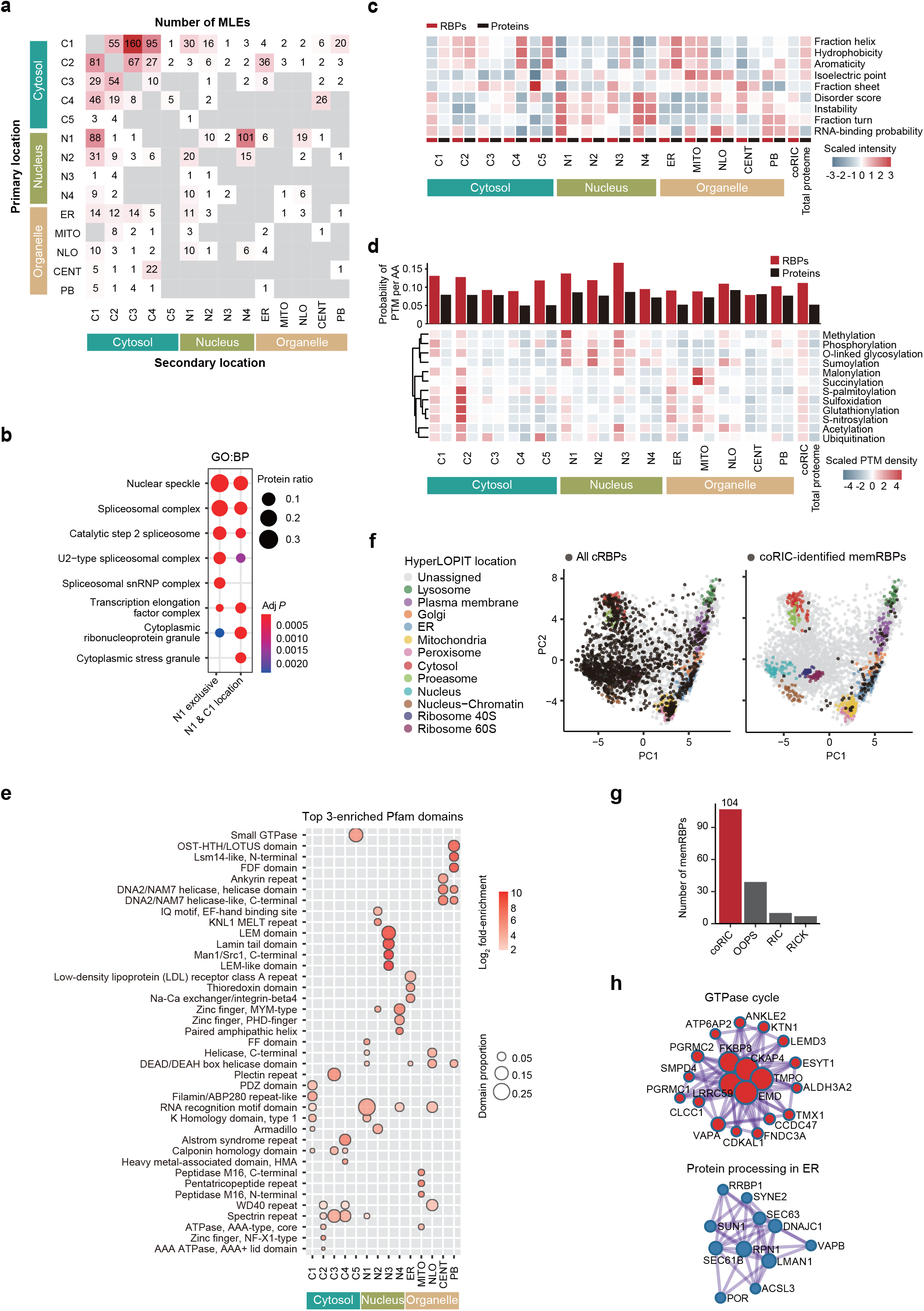
Characteristics of cRBPs (related to Fig. 1). **a,** Heatmap representing the primary and secondary localizations of cRBPs. Proteins with an NMF score of at least 0.15 in each of the two compartments were classified as multi-localized. The intersecting number of MLEs between each pair of compartments is displayed. **b,** GO biological process analysis of the nuclear-exclusively localized cRBPs and nuclear-cytoplasmic localized cRBPs (Fisher’s exact test, Benjamini-Hochberg corrected *P* ≤ 0.01). **c,** Heatmap showing the biophysical properties of the cRBPs and proteins in the corresponding compartments, all cRBPs, and the total proteome (see Materials and Methods). **d,** Bar plot (upper) and heatmap (bottom) showing the sum and individual PTM modification probability per amino acid from different protein groups, respectively. The modification probability of cRBPs and proteins in the corresponding compartment, all cRBPs, and total proteome were calculated and scaled (see Materials and Methods). **e,** Bubble plot showing the top three-enriched Pfam domains in the cRBPs from each compartment (Fisher’s exact test, Benjamini-Hochberg corrected *P* ≤ 0.01). **f,** Projections of all cRBP (left) or coRIC-identified memRBPs (right) to the principal component plot of the HyperLOPIT dataset^22^. Subcellular localization markers from HyperLOPIT are indicated by different colors. PC, principal component. **g,** Bar plot showing the number of memRBPs identified by coRIC, OOPS, RIC, and RICK^10,12,93^. RIC, RNA interactome capture; RICK, capture of the newly transcribed RNA interactome using click chemistry. **h,** Metascape analysis showing the core PPI networks of memRBPs that are enriched in functions related to the GTPase cycle and protein processing in ER. PPI, protein-protein interaction.

**Extended Data Fig. 8.**
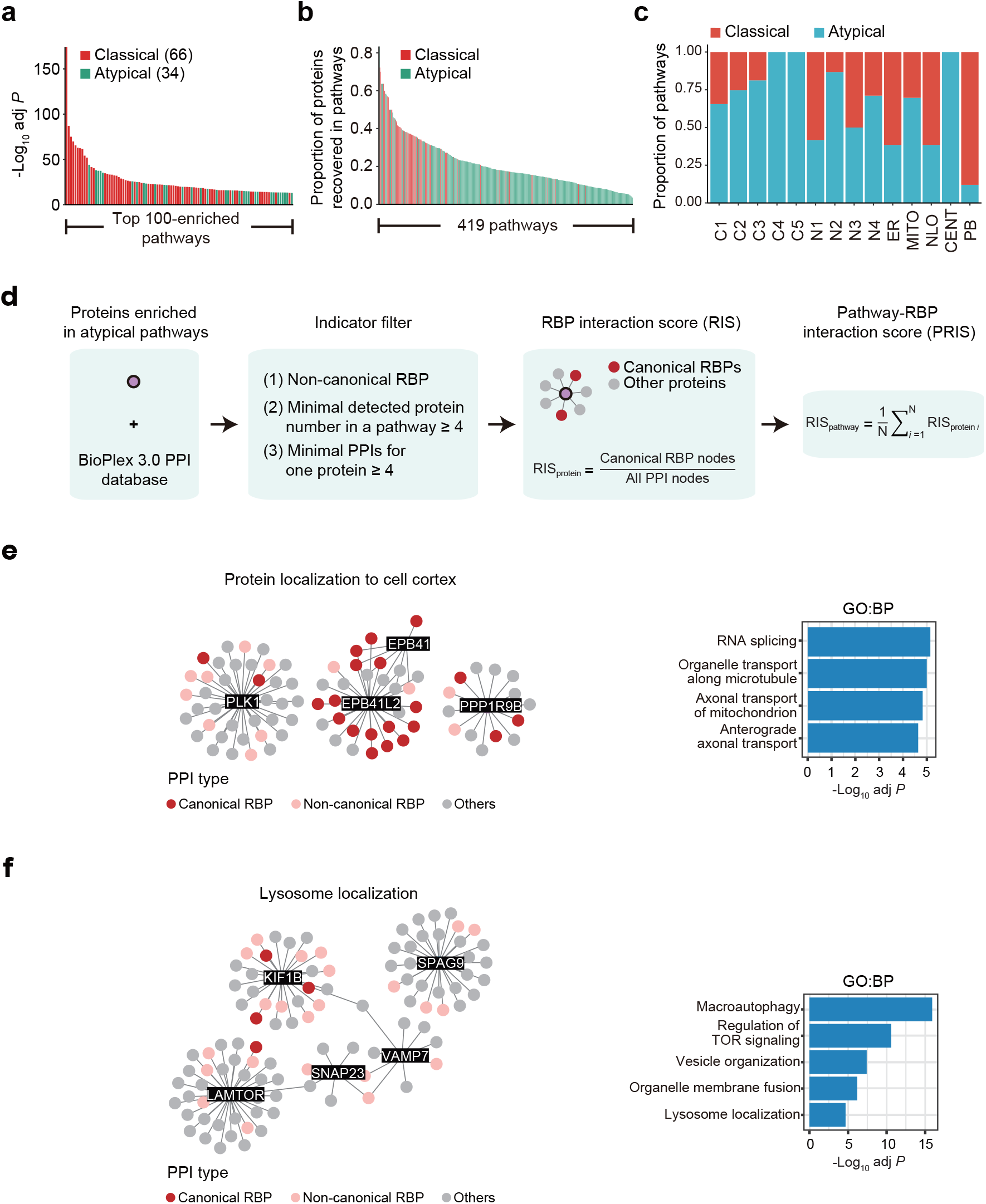
Distribution of cRBPs across biological pathways (related to. Fig. 2**). a,** Bar plot showing the top 100-enriched GO biological process pathways ranked by −log10 adjusted *P* value. Pathways are marked as classical (red) and atypical (green) from the definition in (Fig. 2a) (Fisher’s exact test, Benjamini-Hochberg corrected *P* ≤ 0.01). **b,** Bar plot showing the proportion of proteins recovered in all 419 pathways. Pathways are marked as classical (red) and atypical (green) from the definition in (Fig. 2a). **c,** Bar plot showing the number of classical and atypical pathways enriched in each compartment. **d,** Schematic showing the steps for calculating the pathway-RBP interaction score (PRIS). **(e-f)** PPI networks and PPI-enriched GO biological process terms of the cRBPs in ‘protein localization to cell cortex’ (e) and ‘lysosome localization’ pathway (f) (Fisher’s exact test, Benjamini-Hochberg corrected *P* ≤ 0.01).

**Extended Data Fig. 9.**
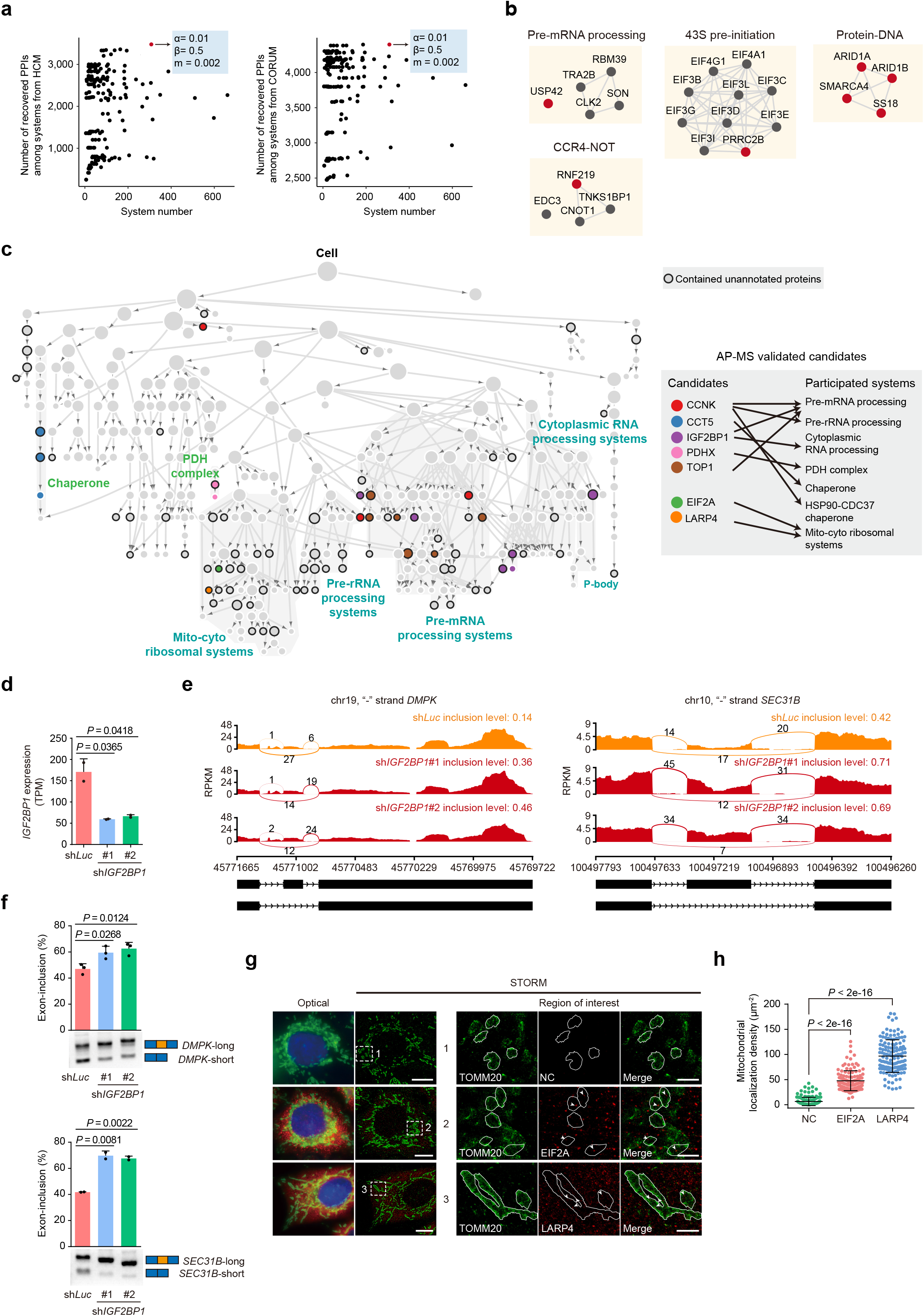
Validation of the RBP hierarchical network (related to Fig. 3). **a,** Dot plots in which each dot is a community hierarchy generated with a particular set of parameters in CliXO (see Materials and Methods). The best performing RBP hierarchical network that recovered most of the PPIs in the two independent datasets HCM (left) and CORUM (right) with the fewest system numbers is highlighted in red. HCM: Human Cell Map. **b,** PPI networks of representative systems from the RBP hierarchical network. Multi-functional cRBPs are colored in red. **c,** Projection of unannotated components and the AP-MS validated candidates into the hierarchical network from (Fig. 3b) (left). Systems containing unannotated components and AP-MS validated candidates are labeled with a black circular border and colored solid circle, respectively. The AP-MS validated candidates and their participated systems are shown (right). **d,** Bar plot showing the *IGF2BP1* expression in the samples from sh*Luc* and sh*IGF2BP1* #1 and sh*IGF2BP1* #2. Data are mean + standard deviation (s.d.) of n = 2 biological replicates. *P* values were generated using a two-tailed Student’s *t*-test. TPM, transcripts per million. **e,** Sashimi plots showing examples of IGF2BP1-modulated skipped exon events. Read counts are from one representative sample of sh*Luc*, sh*IGF2BP1* #1, and sh*IGF2BP1* #2. The inclusion levels are the average of two biological replicates. RPKM, reads per kilobase million. **f,** Semi-quantitative PCR validation of IGF2BP1-modulated skipped exon events. Data are mean + s.d. of n ≥ 2 biological replicates. *P* values were generated using a two-tailed Student’s *t*-test. **g,** Left: Representative conventional images (optical) and STORM renderings of HeLa cells stained for LARP4/EIF2A (red), TOMM20 (mitochondrial membrane, green), and DNA (nuclei, blue). Right: zoom of the regions in the white dotted boxes. The white line indicates the outline of the mitochondria. Arrows indicate typical LARP4 or EIF2A foci localized in the mitochondrial matrix. The negative control (NC) sample was stained with no primary antibody. Scale bars: left, 10 μm; right, 2 μm. **h,** Quantification of the mitochondrial matrix localization density of NC, LARP4, and EIF2A in HeLa cells with STORM. The density was calculated by dividing the mitochondrial localization signal by the area of the mitochondria (μm^2^). Data are the mean ± s.d. of n = 145 mitochondria from 15 cells (NC), n = 123 mitochondria from 9 cells (EIF2A), and n = 134 mitochondria from 9 cells (LARP4). *P* values were generated with a two-tailed Mann-Whitney Wilcoxon test.

**Extended Data Fig. 10.**
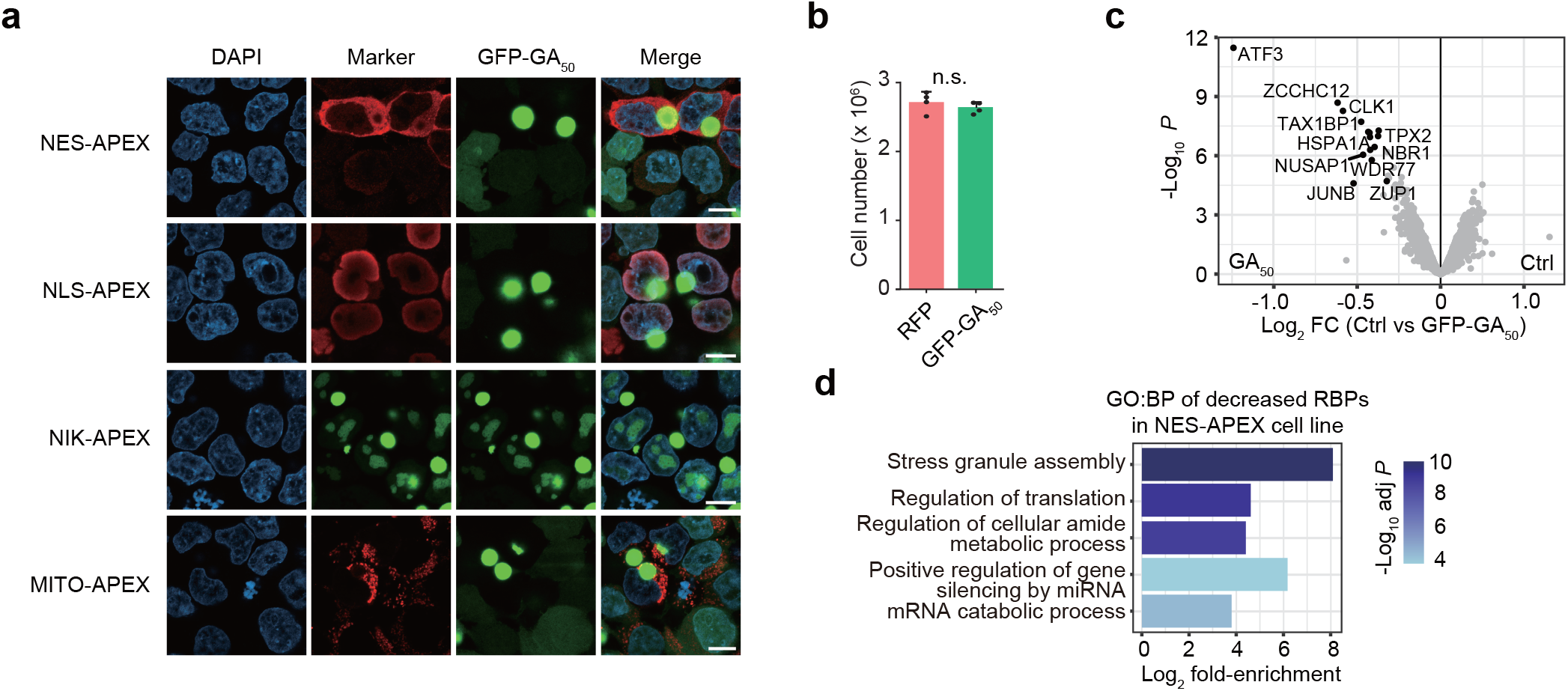
Dynamic changes in RBPs upon polyGA induction (related to Fig. 4). **a,** Confocal immunofluorescence imaging of APEX2 fusion and polyGA localization in four different cell lines. APEX2 fusion protein expression was visualized by V5/FLAG/GFP staining. polyGA was visualized by GFP. DAPI stained nuclei. Scale bars: 10 μm. **b,** Bar plot showing the effect of cell number from RFP and GFP-GA50 induction. Cells were treated with 1μg/mL doxycline for 24 hours and RFP were used as controls. *P* value was generated using a two-tailed Student’s *t*-test. **c,** Volcano plot showing the differentially expressed proteins upon polyGA induction. The log2 FC protein abundance is the average of all biological replicates (moderated t-test, Benjamini-Hochberg corrected P ≤ 0.01). **d,** GO biological process analysis of the proteins with decreased RNA binding upon polyGA induction in the NES-APEX cell line.

**Extended Data Fig. 11.**
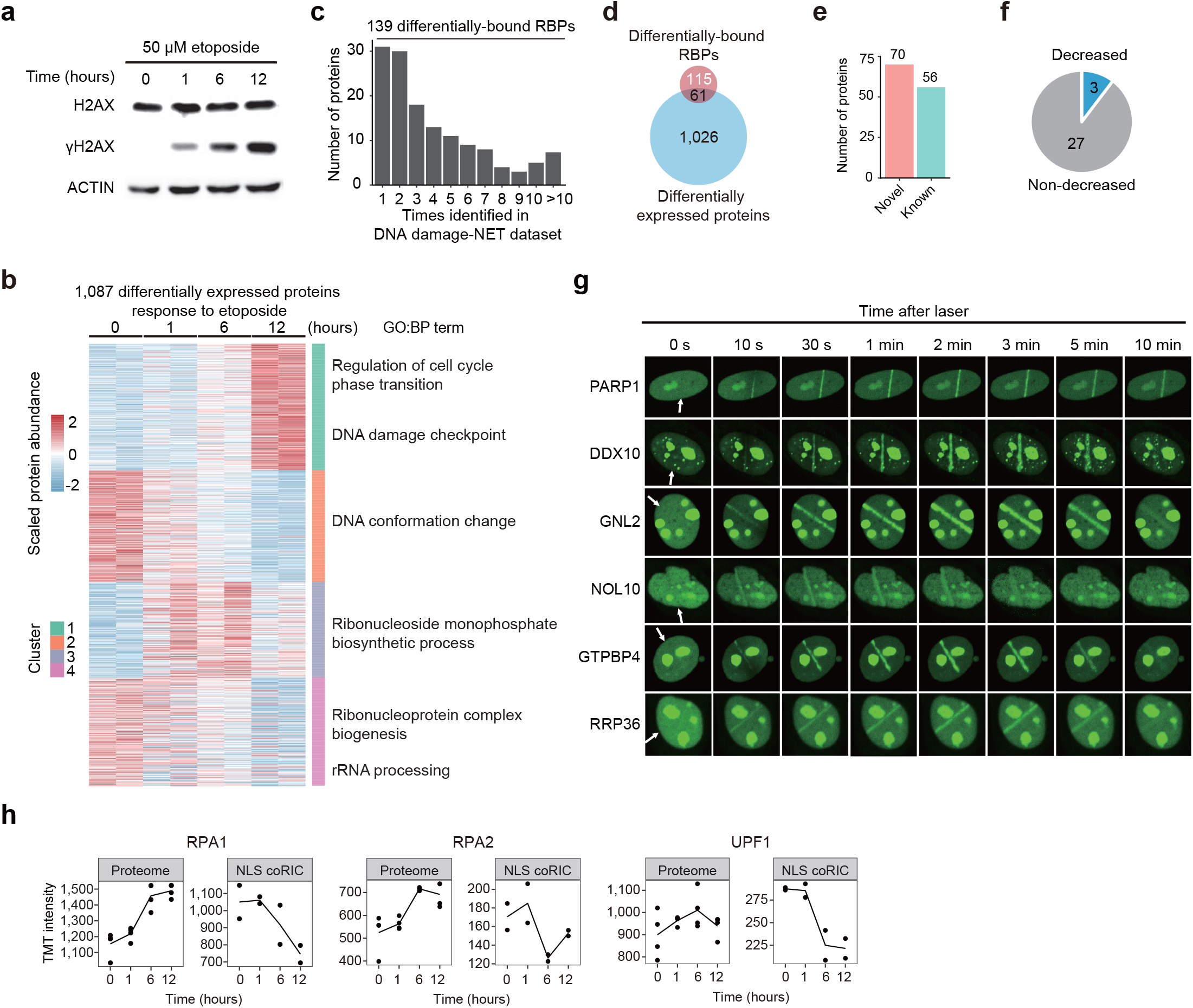
Dynamic changes in RBPs upon etoposide treatment (related to Fig. 4). **a,** Western blot confirming the induction of DNA damage after etoposide treatment at a series of time points (0, 1, 6, and 12 hours). γH2AX is a DNA damage marker. H2AX and ACTIN were the loading controls. **b,** Heatmap showing the 1,087 differentially expressed proteins upon etoposide treatment (moderated *t*-test, Benjamini-Hochberg corrected *P* ≤ 0.01). These proteins were grouped into four clusters based on the scaled protein abundance using hierarchical clustering. Representative GO biological process terms for each group are shown (Fisher’s exact test, Benjamini-Hochberg corrected *P* ≤ 0.01). **c,** Bar plot showing the frequency of occurrence for 139 differentially-bound RBPs induced by etoposide that overlap with the DNA damage-NET dataset^74^. **d,** Venn diagram comparing the 176 differentially-bound RBPs and 1,087 differentially-expressed proteins induced by etoposide. **e,** Bar plot illustrating the number of known and novel RBPs in cluster 1 and 2 from (Fig. 4g) **f,** Pie chart illustrates the protein expression changes of RBP in cluster 3 from (Fig. 4g). **g,** Laser microirradiation assay showing the accumulation of rRNA processing factors at DNA damage loci. White arrows indicate the position of microirradiation. **h,** Line plots showing the protein expression and RNA binding intensity of RPA1, RPA2, and UPF1 upon etoposide treatment. TMT, tandem mass tag.

